# A Non-Canonical Interface of SNX9 PX Domain Selectively Sequesters PI(3,4)P_2_ Lipids, Protecting Them from Hydrolysis

**DOI:** 10.1101/2025.03.26.645564

**Authors:** Jeriann R. Beiter, Chieh-Ju Sung, Shan-Shan Lin, Nicolas de Vuono, Senthil Arumugam, John Manzi, Aurélie Bertin, Patricia Bassereau, Ya-Wen Liu, Gregory A. Voth, Feng-Ching Tsai

**Affiliations:** Department of Chemistry, Chicago Center for Theoretical Chemistry, Institute for Biophysical Dynamics, and James Franck Institute, The University of Chicago, Chicago, IL 60637, USA; National Taiwan University College of Medicine, Institute of Molecular Medicine, TW; Institut Curie, Université PSL, Sorbonne Université, CNRS UMR168, Physique des Cellules et Cancer, 75005 Paris, FR; European Molecular Biological Laboratory Australia (EMBL Australia) and Monash Biomedicine Discovery Institute, Faculty of Medicine, Nursing and Health Sciences, Monash University, Clayton/Melbourne, AU

## Abstract

Plasma membrane remodeling processes are tightly regulated by the spatiotemporal distribution and dynamic conversion of phosphoinositidyl lipids (PIPs). This regulation is controlled by the recruitment of proteins such as sorting nexin 9 (SNX9), a key mediator of late-stage endocytosis and macropinocytosis. Using live cell imaging, *in vitro* reconstitution assays, and molecular dynamics simulations, we investigated how SNX9 distinguishes between PI(3,4)P_2_ and PI(4,5)P_2_, and the physiological relevance of this selectivity. Our results revealed that during macropinocytic membrane ruffling, SNX9 is recruited in a spatiotemporally coordinated manner with PI(3,4)P_2_, but not with PI(4,5)P_2_. While SNX9 induces comparably weak mechanical remodeling on model membranes containing either PIP_2_ species, it exhibits a clear selective binding to PI(3,4)P_2_, mediated by a non-canonical interface. Through mutational analysis of key residues involved in this sequestration, we further demonstrated that SNX9 protects PI(3,4)P_2_ from hydrolysis. Together, these results reveal a previously unrecognized mechanism of SNX9-PIP_2_ lipid interaction that underscores SNX9’s pivotal role in coordinating membrane remodeling processes.

**Teaser:** Curvature sensing BAR protein SNX9 selectively sequesters PI(3,4)P_2_ lipids, acting as a checkpoint in cell membrane remodeling.

## Introduction

Cells rely on the dynamic and localized synthesis of phosphoinositide (PIP) lipids to generate specialized membrane regions that recruit PIP-associated peripheral proteins at precise locations and times. This intricate PIP synthesis regulation enables numerous membrane processes, especially in endocytic pathways, such as clathrin-mediated and clathrin-independent endocytosis, macropinocytosis, phagocytosis, and pinocytosis. During these endocytic processes, kinases and phosphatases catalyze PIPs to produce different PIP species sequentially. For example, in macropinocytosis, the enrichment of PIP lipids proceeds from PI(4,5)P_2_ to PI(3,4,5)P_3_, then PI(3,4)P_2_, and finally PI(3)P, a marker of early endosomal membranes (*1–3*). Similarly, clathrin- mediated endocytosis begins with PI(4,5)P_2_ enrichment, followed by PI(3,4)P_2_, and then PI3P (*4, 5*). PI(4,5)P_2_ is known to be abundant at the plasma membrane, where its local concentration often increases during endocytosis. In contrast, other PIPs like PI(3,4)P_2_ are synthesized transiently and at much lower concentrations at endocytic sites, with PI(3,4)P_2_ level being ∼ 40- fold lower than PI(4,5)P_2_ (*6*). To recognize and bind specific PIP lipids, many membrane- associated proteins have PIP-binding domains such as PX (<u>p</u>hox <u>h</u>omology), PH (<u>p</u>leckstrin <u>h</u>omology), FYVE (<u>F</u>ab1, <u>Y</u>OTB, <u>V</u>ac1 and <u>E</u>EA1), and C2 domains (*7*). These domains often display affinity for multiple PIP species. For instance, various PX domains bind both PI(4,5)P_2_ and PI(3,4)P_2_, though with different affinities (*8*). Discriminating between similar PIPs, such as PI(3,4)P_2_ and PI(4,5)P_2_, is challenging due to their nearly identical headgroups, which share the same charges, sizes, and protonation constants (Fig. 1A) (*9, 10*). This overlapping specificity of similar PIPs raises fundamental questions: How do protein domains achieve selective recognition of specific PIP lipids? How might PIP-binding proteins impact PIP conversions, possibly by affecting phosphatase accessibility and thus modulating endocytic processes? Addressing these questions is fundamental for deciphering the regulation of membrane dynamics in cells.

**Fig. 1.**
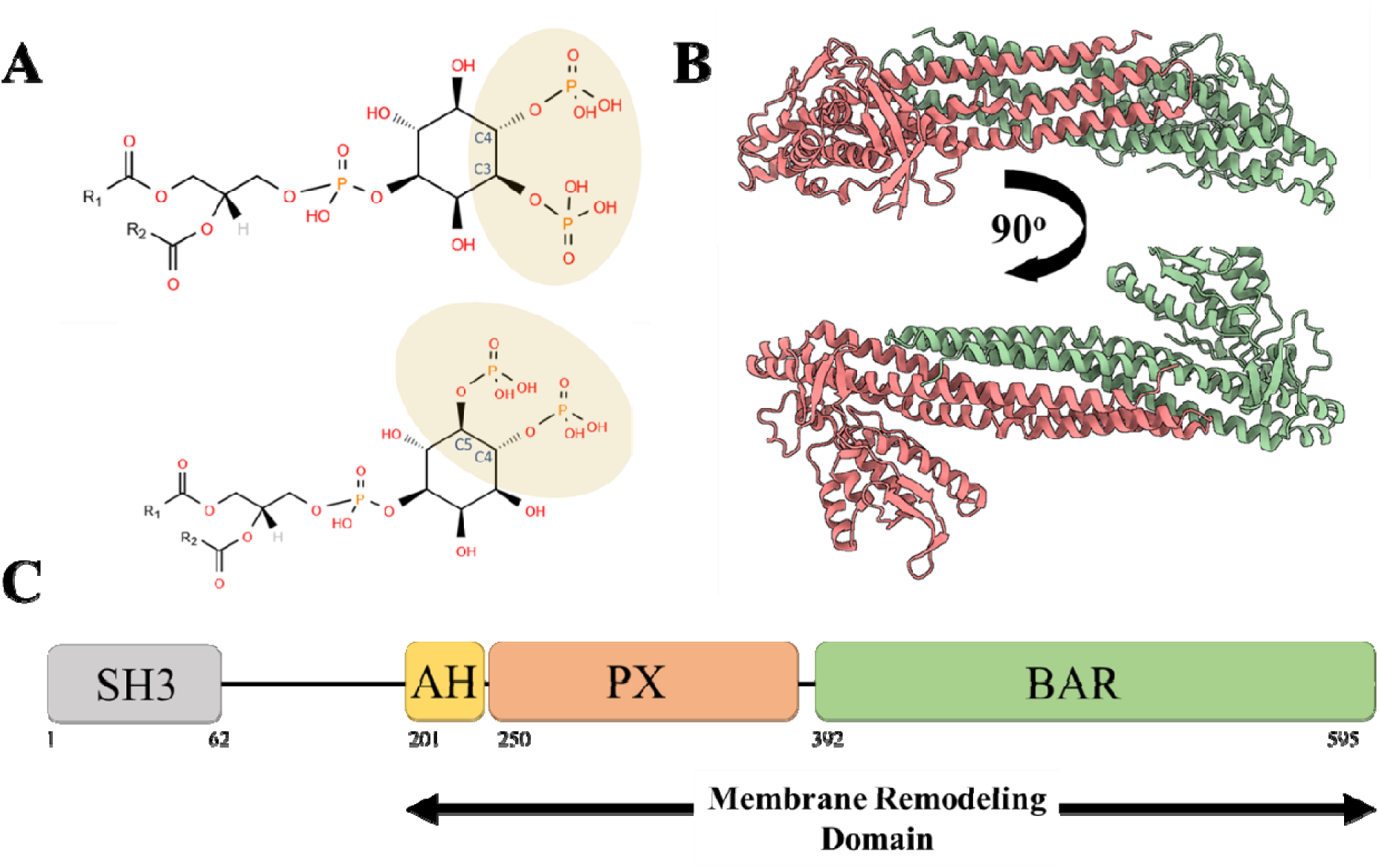
Structural information of sorting nexin 9 (SNX9), PI(3,4)P_2_, and PI(4,5)P_2_. **(A)** Stereoprojected line structures of the headgroups of PI(3,4)P_2_ (*Top*) and PI(4,5)P_2_ (*Bottom*), with the C3 and C4 or C4 and C5 inositol carbons (respectively) labelled to demonstrate the location of the phosphate functional groups. Line structures obtained from LIPID MAPS® structure database. **(B)** Predicted structure of the membrane remodeling domain of SNX9 dimer viewed from the side (*Top*) and looking top-down (*Bottom*). The predicted structure has an RMSD from that of the apo crystal structure of 1.196 Å (PDB ID: 2RAI) and of the PI(3)P bound crystal structure of 1.271 Å (PDB ID: 2RAK). **(C)** Schematic of domains of SNX9, in sequence order. The range from residues 200-595 is collectively known as the membrane remodeling domain. SH3 – SRC Homology 3; AH – amphipathic helix; PX – Phox homology; BAR – Bin/Amphiphysin/Rvs domain.

The crystal structures of PIP-associated domains typically reveal a single PIP bound in a canonical binding pockets (*11–14*). However, there is growing evidence that many protein domains interact with lipids in a promiscuous, multivalent manner (*15, 16*). Superstoichiometric lipid binding, where one protein domain can simultaneously bind multiple lipids of the same species, often involves additional secondary binding pockets or polybasic patches and motifs that promote such interactions (*16–18*). For instance, recent molecular dynamics (MD) simulations have shown that domains like the FERM domain of ezrin can bind simultaneously to multiple PI(4,5)P_2_ lipids, and the PH domain of GRP1, with at least two PIP_3_ lipids (*19–21*). However, concurrent with superstoichiometric binding, many peripheral membrane proteins contain PH and PX domains that are relatively weak binders yet still exhibit certain PIP selectivity (*8, 22*). How these PIP-binding domains achieve superstoichiometric yet selective PIP binding remains an open question in understanding the molecular mechanisms of lipid-protein interactions.

In this study, to reveal how certain proteins achieve both superstoichiometric and selective PIP binding, we investigated how the PX-BAR domain of sorting nexin 9 (SNX9) interacts with PI(4,5)P_2_ and PI(3,4)P_2_, two key PIP lipids in SNX9-mediated endocytosis and macropinocytosis (*23, 24*). SNX9 consists of an N-terminal Src Homology 3 (SH3) domain, a long unstructured motif termed the “low complexity” (LC) domain, and a C-terminal PX-BAR unit, which comprises a PX domain paired with a <u>B</u>in/<u>A</u>mphiphysin/<u>R</u>vs (BAR) domain (Fig. 1, B and C). BAR domains are known to bind to negatively charged lipids, including PIPs and phosphoserine (PS) lipids; moreover, many BAR domains have been shown to be able to sense and generate curved membranes (*25*). SNX9 has been shown to play a prominent role in clathrin-mediated endocytosis and macropinocytosis, by facilitating plasma membrane deformation and by recruiting partners such as dynamin and actin nucleation promoting factor N-WASP (*24, 26, 27*). While SNX9’s PX domain is thought to assist in targeting PIP-rich membranes, its PIP binding selectivity remains unclear (*23*). Many studies have investigated SNX9’s PIP preference: some indicate PI(3,4)P_2_ and PI(4,5)P_2_ as predominant, while others suggest a preference for PI3K products such as PI(3)P, PI(3,4)P_2_, and PIP_3_ (*8, 28, 29*). Other work indicates that SNX9 binds both PI(3,4)P_2_ and PI(4,5)P_2_ weakly, and is recruited to membranes through a general electrostatic mechanism (*30*). While the high resolution SNX9 PX-BAR crystal structure indicates the binding of the short-chain PI(3)P to the canonical binding pocket of the PX domain, it does not clarify selective binding mechanism for other PIP_2_ lipids (PDB ID: 2RAK (*11*)). Unlike PI(3)P, which has a smaller charged headgroup, the more complex PIP_2_ headgroups suggest that electrostatic and steric factors might allow the PX-BAR domain to achieve specific PIP binding, particularly with the additional consideration of superstoichiometric lipid binding.

In this study, we conducted live-cell imaging to quantify the temporal recruitment of SNX9 to macropinocytic membrane ruffles in relation to PIP enrichment. We observed that SNX9 is recruited synchronously with PI(3,4)P_2_ but is delayed relative to PI(4,5)P_2_. To reveal how SNX9 distinguishes between PI(3,4)P_2_ and PI(4,5)P_2_, we performed *in vitro* reconstituted assays. We found that SNX9 binds both PIPs with comparable affinities, and generates and senses membrane curvature irrespective of the two PIP_2_ species. Motivated by these findings, we next examined how SNX9 modulates PI(3,4)P_2_ and PI(4,5)P_2_ dynamics at the molecular scale. Atomistic MD simulations of SNX9 PX-BAR domain with model membranes revealed a non-canonical binding interface in the PX domain that selectively favors PI(3,4)P_2_ over PI(4,5)P_2_, and drives PI(3,4)P_2_ sequestration. We hypothesized that this selective sequestration may influence the conversion of PI(3,4)P_2_ during macropinocytosis. Indeed, mutational analysis in cells confirmed that this non- canonical interface can interfere with the downstream phosphatase INPP4B’s ability to convert PI(3,4)P_2_ into PI3P. Collectively, our findings suggest that the selectively sequestration of PI(3,4)P_2_ by SNX9 may not only facilitate its recruitment to the plasma membrane, but also provides a “protective” effect, potentially preventing PI(3,4)P_2_ from being prematurely converted into PI(3)P during macropinocytosis and endocytosis.

## Results

### SNX9’s Recruitment to Ruffling Membranes is Coincident with PI(3,4)P_2_ but Not PI(4,5)P_2_

To investigate the interplay between SNX9 and PI(4,5)P_2_ or PI(3,4)P_2_ during macropinocytosis, we conducted live-cell imaging to monitor their spatiotemporal distribution upon stimulation with platelet-derived growth factors (PDGF) in serum-starved cells. The initial stage of macropinocytosis involves membrane ruffle formation, a process where SNX9 plays a crucial role (*31*). We performed time-lapse imaging of cells overexpressing fluorescently tagged SNX9 alongside fluorescent PH-domain probes for PI(4,5)P_2_ (PLCδ-PH-EGFP) or PI(3,4)P_2_ (NES- EGFP-cPHx3) to track the recruitment and dynamics of each molecule at the sites of ruffle formation (Fig. 2, A and C). Our results show that SNX9 co-localizes spatially with both PIP_2_ reporters at PDGF-induced membrane ruffles (Fig 2, B and D). However, quantitative analysis of SNX9 and PIP_2_ fluorescence intensity profiles during the first 60 seconds of ruffle formation shows distinct enrichment kinetics. Specifically, for PI(4,5)P_2_, we observe a delayed appearance of SNX9 fluorescence relative to PI(4,5)P_2_ at the same membrane ruffle site (Fig. 2E). In contrast, SNX9 and PI(3,4)P_2_ fluorescence signals appear simultaneously, with no detectable lag (Fig. 2F). The observed delayed, yet spatially colocalized, appearance of SNX9 relative to PI(4,5)P_2_ is consistent with the established role of PI(4,5)P_2_ as an upstream factor during endocytosis and macropinocytosis (*1*). However, the simultaneous appearance of SNX9 with PI(3,4)P_2_, as opposed to the lag observed with PI(4,5)P_2_, suggests a potential selective association between SNX9 and PI(3,4)P_2_ at macropinocytic ruffles. Our finding suggests that SNX9 displays a specificity for PI(3,4)P_2_ over PI(4,5)P_2_. This observation is consistent with previous studies indicating that PI(3,4)P_2_ promotes SNX9 membrane association at the late stage of clathrin-mediated endocytosis, thereby facilitating dynamin recruitment by SNX9 to drive membrane fission (*30, 32*).

**Fig. 2.**
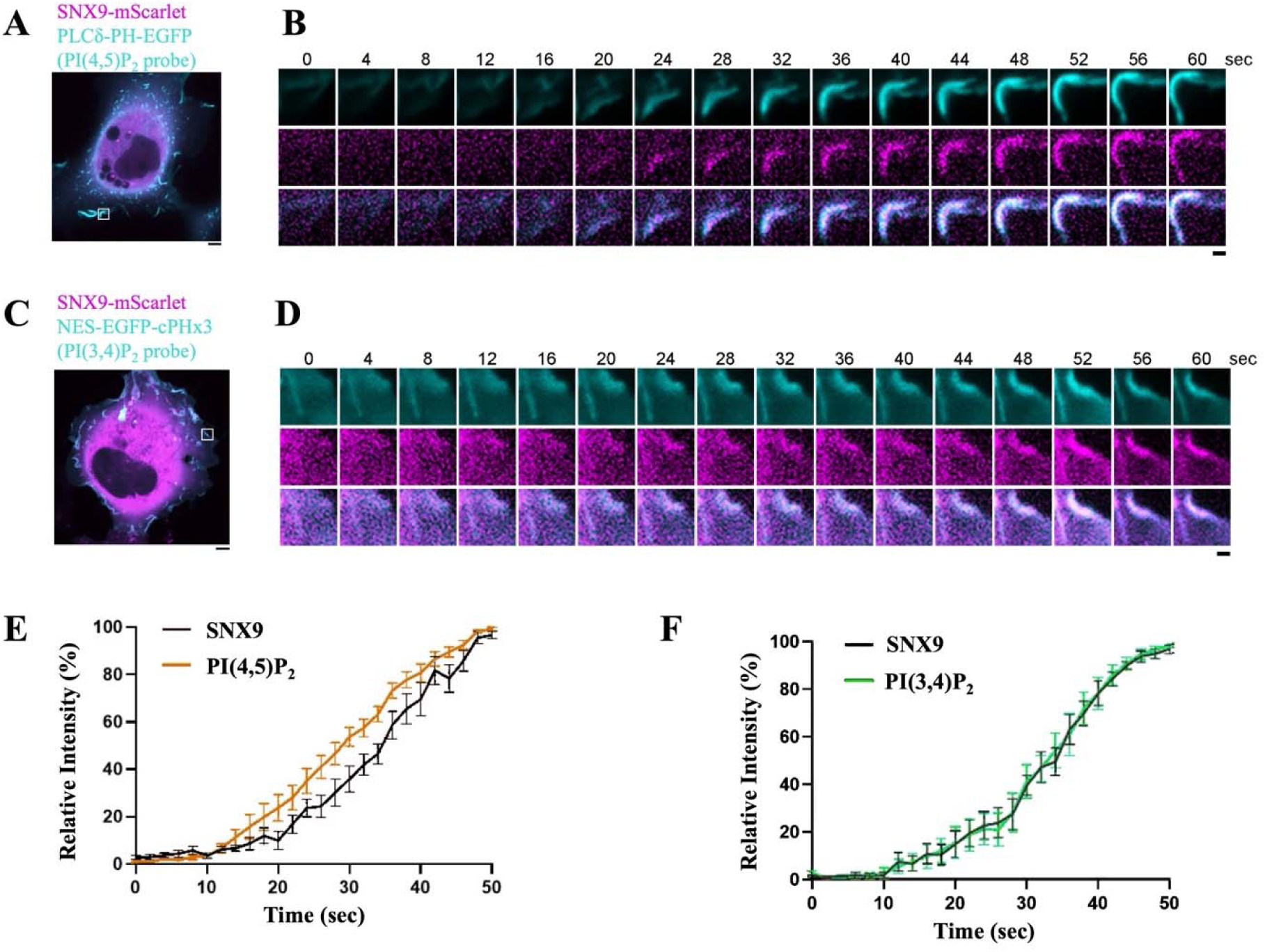
SNX9 recruitment to macropinocytic ruffles is coincident with PI(3,4)P_2_ rather than PI(4,5)P_2_. **(A)** Representative overlay image of SNX9 (magenta) and PI(4,5)P_2_ fluorescent probe (cyan) signals. The white inset box highlights the ruffle shown in (B). **(B)** Time series evolution of the signals of SNX9 (magenta) and PI(4,5)P_2_ probe (cyan) from a macropinocytic ruffle over the first 60 seconds of ruffle formation. **(C)** Representative overlay image of SNX9 (magenta) and PI(3,4)P_2_ fluorescent probe (cyan) signals. The white inset box highlights the ruffle shown in (D). **(D)** Time series evolution of the signal of SNX9 (magenta) and PI(3,4)P_2_ probe (cyan) from a macropinocytic ruffle over the first 60 seconds of ruffle formation. **(E)** Relative intensity of fluorescent signals for PI(4,5)P_2_ reporter and SNX9 from time series, quantified over 12 ruffles. **(F)** Relative intensity of fluorescent signals for PI(3,4)P_2_ and SNX9 from time series, quantified over 11 ruffles. Data represent mean ± standard error of the mean. Scale bars, 5 μm (A and C), 1 μm (B and D).

### SNX9 is a Weakly Scaffolding BAR Protein that Senses and Generates a Wide Range of Membrane Curvatures on both PI(3,4)P_2_- and PI(4,5)P_2_-Containing Membranes

Given that SNX9 is a BAR domain protein, we next asked how different PIP_2_ species in membranes impacts the expected ability of SNX9 to sense and generate curved membranes. We purified full length SNX9 and labelled it with Alexa fluorophore dyes for detection by confocal microscopy. We first determined whether SNX9’s binding affinity to flat membranes differs between PI(4,5)P_2_ and PI(3,4)P_2_. To do this, we used giant unilamellar vesicles (GUVs) with diameters of about 5 μm or larger. GUV membranes were composed of eggPC, supplemented with 10 mol% DOPS, 10 mol% DOPE, 15 mol% cholesterol, and 0.5 mol% Bodipy ceramide, along with either 8 mol% PI(4,5)P_2_ or PI(3,4)P_2_. Upon incubating SNX9 with GUVs, we observed the formation of outward membrane tubules on the GUVs, confirming SNX9’s ability to deform membranes and generate membrane tubules, a characteristic feature of BAR domains (Fig 3, A and B). To estimate the binding affinities of SNX9 on PI(4,5)P_2_- and PI(3,4)P_2_-containing GUVs, we measured the surface density of SNX9 on GUV membranes as a function of its bulk concentration (Fig. 3, C and D) (*33*). By fitting to the Hill equation, we estimated dissociation constants of 31 nM for PI(4,5)P_2_ and 47 nM for PI(3,4)P_2_. These comparable affinities are consistent with previous findings for the PX-BAR domain of SNX9 on small unilamellar vesicles (*30*).

**Fig. 3.**
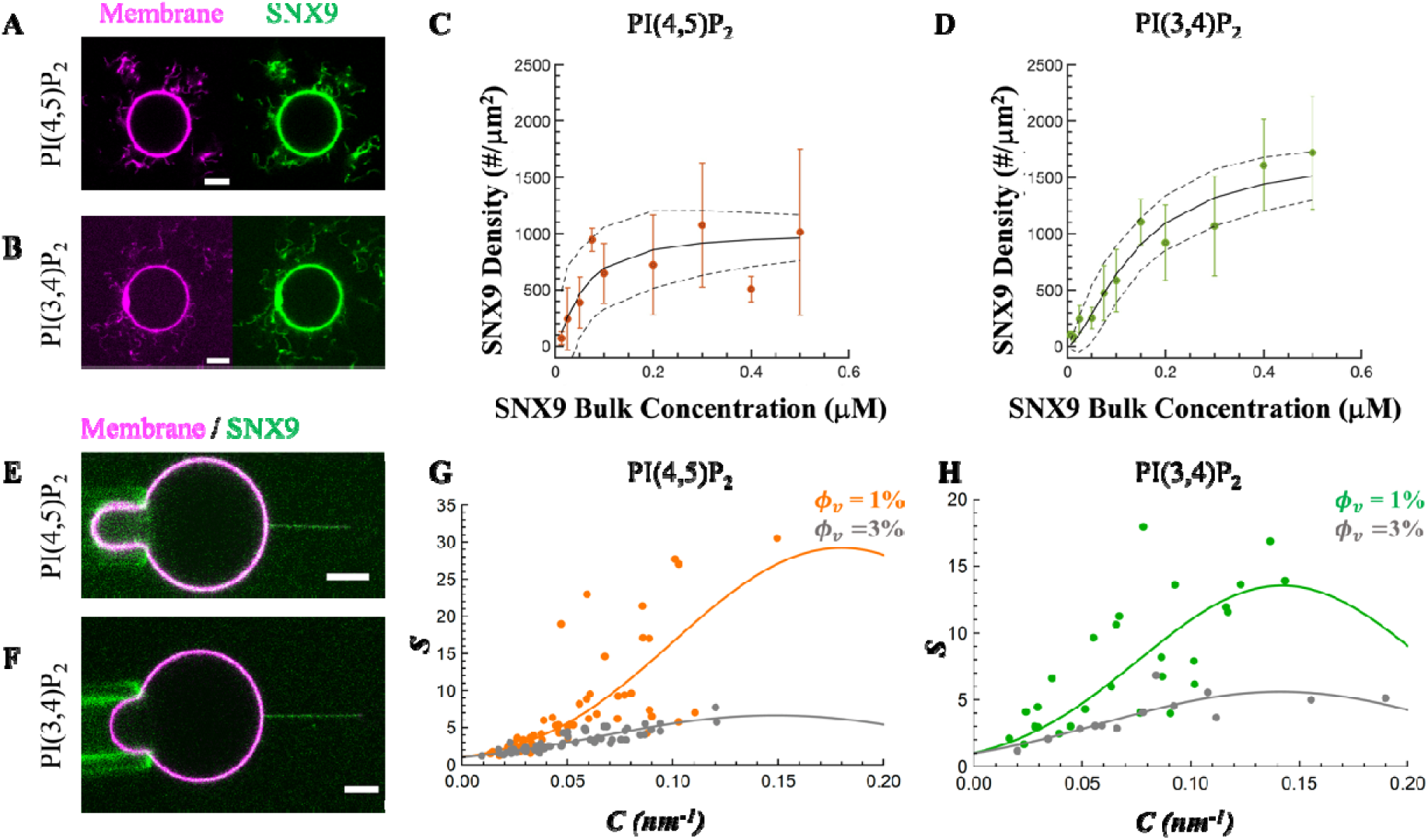
SXN9 generates membrane tubes and senses membrane curvature with comparable affinities on PI(3,4)P_2_ and PI(4,5)P_2_-GUVs. **(A and B)** GUVs exhibiting outward tubes generated by SNX9. Green, SNX9 and Magenta, membranes. **(C and D)** Analysis of SNX9’s binding on GUVs. Data fitted to the Hill equation (Equation. 1, Materials and Methods) to obtain the dissociation constant and the Hill coefficient n=1.18 for PI(4,5)P_2_ GUVs and and n=1.55 for PI(3,4)P_2_ GUVs. Black lines represent the fitted regression model, and dashed black lines, the bounds of the 95% confidence interval. For each bulk concentration, PI(4,5)P_2_, N = 4, 4, 4, 1, 4, 2, 3, 1, 3 independent experiments, n = 81, 78, 70, 32, 73, 61, 68, 21, 41 GUVs; PI(3,4)P_2_, N = 1, 2, 2, 2, 3, 3, 1, 3, 3, 1, 1 independent experiments, n = 30, 52, 64, 55, 79, 72, 34, 80, 75, 30, 28 GUVs. **(E and F)** Representative confocal image of GUVs with pulled tethers. Tube radius ∼11 nm (E) and ∼ 9 nm (F). **(G and H)** Sorting ratio as a function of tube curvature, *C=1/R*. Protein coverage on GUVs,, 1% (orange for PI(4,5)P_2_ and green for P(3,4)P_2_) and 3% (grey for both PIP_2_). By fitting the sorting curves to Equation 3, we obtained averaged 1/ 6 nm and 7 nm for PI(4,5)P_2_ and PI(3,4)P_2_, respectively, and averaged 4 and 5 for PI(4,5)P_2_ and PI(3,4)P_2_, respectively. N = 8 and 2 independent experiments, n = 21 and 9 GUVs, for PI(4,5)P_2_ and PI(3,4)P_2_, respectively. All scale bars, 5 μm.

To evaluate SNX9’s ability to sense membrane curvature, we generated cylindrical membrane nanotubes from SNX9-coated GUVs using optical tweezers, with tube radii that can be tuned by changing membrane tension via micropipette aspiration (*33*) (Fig. 3, E and F). We quantified the enrichment of SNX9 on the nanotubes relative to the flat GUV membranes by calculating the sorting ratio, as previously described for other BAR domains (*33*). Our data show that SNX9’s sorting ratio increases as the membrane tubes are thinner – that is, as curvature increases - within the experimentally accessible range of ∼ 0.0125 nm^-1^ to ∼ 0.15 nm^-1^ (corresponding to tubes radii of ∼ 80 nm and 7 nm, respectively) (Fig. 3, G and H). Furthermore, by comparing the sorting ratio of SNX9 at a relatively low surface density (on average 1% surface coverage, i.e. protein areal density 200 μm^-2^), where enrichment effects are expected to be the most pronounced, we observed that SNX9 are enriched on both PI(4,5)P_2_ and PI(3,4)P_2_ membrane tubes, as indicated by the sorting ratios larger than 1 (Fig. 3, G and H). Because the full-length SNX9 contains a PX- BAR domain, to analyze its sorting behavior, we applied the curvature mismatch model (*34*), which has been shown to be more appropriate for full length BAR protein amphiphysin, and truncated BAR-PH domain of β2-centaurin than the spontaneous curvature model (*35*). This analysis enables us to estimate the spontaneous curvature of membrane-bound SNX9 (*C̅_p_*) and the associated elastic constant *κ̄* - which reflects the energetic cost of membrane deformation and indicates the strength of SNX9’s mechanical capacity to induce curvature. Our analysis indicates that the intrinsic curvatures of membrane-bound SNX9 for PI(4,5)P_2_ and PI(3,4)P_2_ containing membranes are comparable (1/*C̅_p_* ∼ 6 nm and ∼ 7 nm, respectively), as are their elastic constants (*κ̄* ∼ 4 kBT and ∼5 k_B_T, respectively). These findings suggest that, at a macroscopic level, SNX9 has a similar sorting capacity and mechanical effect on membranes, regardless of the PIP_2_ species.

Moreover, the relatively low elastic constants imply that SNX9 is a weaker membrane scaffolding protein as compared to other BAR proteins such as amphiphysin, β2-centaurin, and IRSp53 (*35*).

To gain molecular scale insights into how SNX9 organizes and deforms PI(4,5)P_2_- and PI(3,4)P_2_- containing membranes, we performed cryo-electron microscopy (cryo-EM). A heterogeneous solution of Large Unilamellar Vesicles (LUVs) with diameters ranging from 50 nm to 1 µm was prepared by resuspending dry lipid films composed of DOPC and 15% DOPS, supplemented with 10% PI(4,5)P_2_ or PI(3,4)P_2_. Besides spherical vesicles, this protocol also yields tubulated vesicles in roughly 15% of the vesicles (*36*). These vesicles were incubated with 1 μM SNX9 for at least 1 hour at room temperature before being plunge frozen onto EM grids. Our cryo-EM results revealed dense SNX9 binding along membrane tubes under both PIP_2_ conditions, whereas the remainder of the liposome membranes exhibited a lower density of SNX9 (Figs. 4, A and B, *Left* panel). Notably, unlike previous studies where BAR domains, such as endophilin, displayed a well-registered alignment on membrane tubes, SNX9 exhibits a more disorganized binding pattern (Fig. 4, A and B, *Left* panel, arrows) (*37*). This less ordered organization likely reflects the influence of the PX domain and disordered regions present in full-length SNX9. Although we attempted to purify the SNX9 BAR domain to further test this hypothesis, the isolated domain proved too unstable for performing experiments (*11*).

**Fig. 4.**
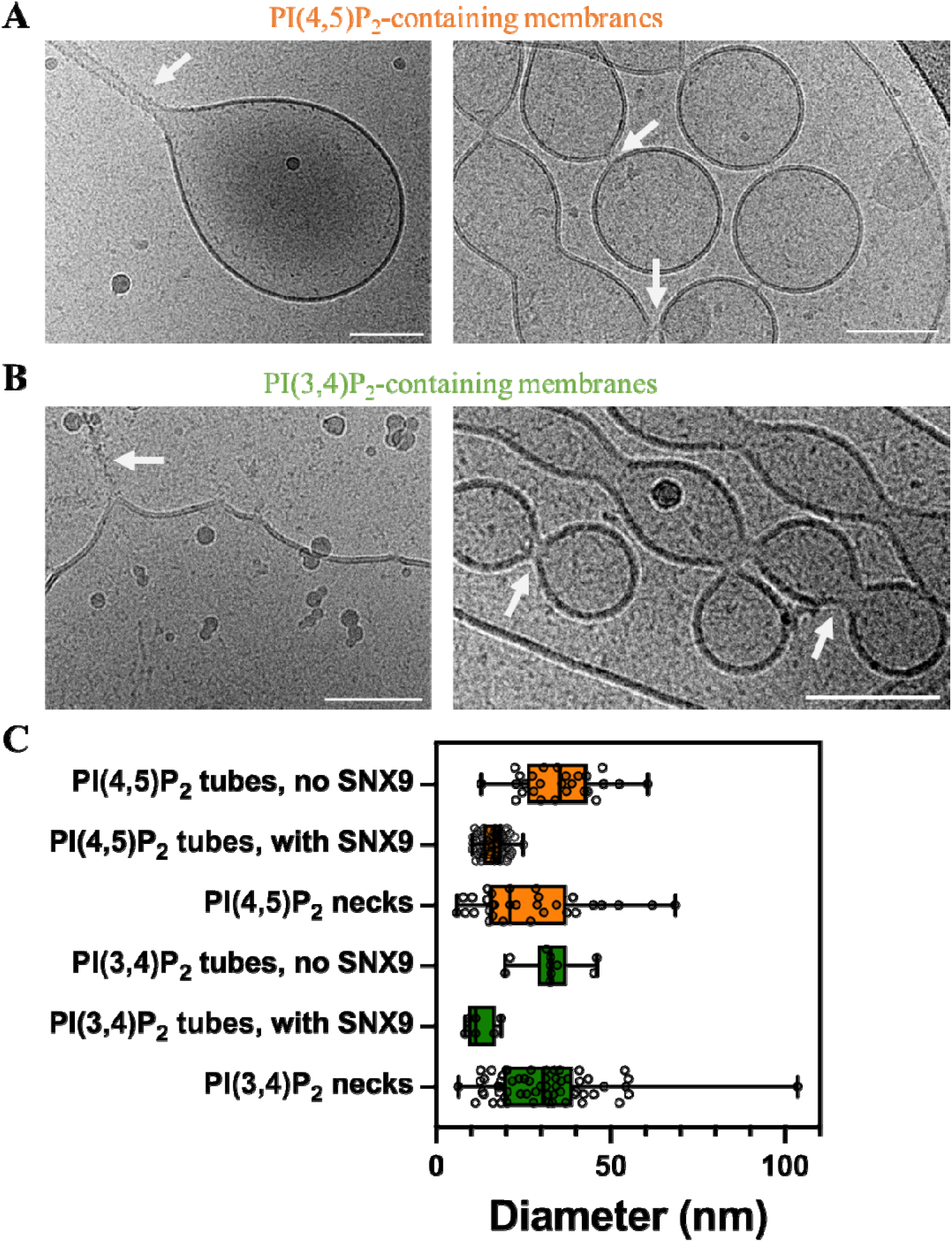
SNX9 induces remodeling of membranes containing either PI(4,5)P_2_ or PI(3,4)P_2_. **(A and B)** Representative cryo-EM images of full length SNX9 incubated with membranes containing PI(4,5)P_2_ or PI(3,4)P_2_, and with tube and neck-like structures indicated by white arrows. Scale bars, 100 nm. **(C)** Quantification of the diameters of observed membrane structures, including tubes with no apparent SNX9 signals, tubes with apparent SNX9 signals, and neck-like structures with apparent SNX9 signals. n = 28, 68, 31, 10, 6, 60 structures analyzed from data presented *Top* to *Bottom*.

Quantitative analysis of tube diameters revealed that SNX9-decorated tubes had median diameters of 16.7 nm for PI(4,5)P_2_- containing membranes and 11.4 nm for PI(3,4)P_2_-containing membranes. In contrast, bare membranes exhibited significantly larger medians of 35.5 nm for PI(4,5)P_2_ and 32.9 nm for PI(3,4)P_2_ (Fig. 4C). These data confirm that SNX9 can deform membranes and constrict tubes to diameters near 15 nm, consistent with the spontaneous curvature measured in our tube pulling experiments. We noted that to compare the intrinsic curvatures obtained by tube pulling experiments with those from Cryo-EM, a half-bilayer thickness (2 nm considering tube radius and 4 nm, diameter) should be added, resulting in a tube diameter of 16 nm and 18 nm for PI(4,5)P_2_ and for PI(3,4)P_2_, respectively. For both PIP_2_ conditions, we observed peanut-shaped vesicles characterized by a narrow neck-like region at their center. At these necks, the membranes appeared less defined and more diffuse than in the remainder of the vesicles, suggesting local SNX9 enrichment at these necks and reinforcing its role in membrane remodeling (Fig. 4, A and B, *Right* panel). Notably, these neck regions exhibited a broader diameter distribution compared to the SNX9-decorated tubes, indicating that SNX9 functions as a weak scaffolding protein capable of accommodating a wide range of membrane curvatures (Fig. 4C). This observation aligns with our tube pulling experiments, where we observed a wide distribution of sorting ratios at given membrane curvatures (Fig. 3, G and H). Of note, the stability of these neck-like structures was confirmed by overnight incubation experiments. Together, our Cryo-EM findings demonstrate that SNX9 preferentially associated with curved, tubular membranes in both PI(4,5)P_2_ and PI(3,4)P_2_ membranes. Moreover, SNX9 not only binds loosely to membrane tubes but also induces the formation of membrane necks, progressively transforming vesicles into peanut-shaped structures with pinched necks and ultimately into thin tubes.

### SNX9 PX-BAR Drives Distinct Distributions of PI(3,4)P_2_ and PI(4,5)P_2_ Lipids

To explore the molecular mechanisms by which SNX9 interacts with membranes containing PI(4,5)P_2_ and PI(3,4)P_2_, we conducted all-atom molecular dynamics (AA-MD) simulations of the dimeric SNX9 membrane remodeling region, including the PX-BAR domain (residues 200-595) with model membranes containing the following compositions: DOPC:DOPS 80:20; DOPC:DOPS:PI(3,4)P_2_ 80:15:5; and DOPC:DOPS:PI(4,5)P_2_ 80:15:5 (Fig. 5A). The structure of the membrane remodeling region dimer was predicted using AlphaFold2, to facilitate the realistic inclusion of the C-terminal amphipathic helix and missing loops (*38*). By probing the unbiased interactions between SNX9 PX-BAR and model membranes, we aimed to reveal whether there is a difference in the way that SNX9 interacts with PI(3,4)P_2_ and PI(4,5)P_2_. Specifically, we sought to investigate how SNX9 PX-BAR influences the local dynamics of PIP_2_ lipids upon binding. Although the timescales achievable by AA-MD (a total of 5 microseconds for each composition) limits direct measurement of kinetic rate constants, the simulations enable detailed analysis of nanoscopic phenomena such as lipid diffusion. To this end, we calculated the total distance traversed by all lipids and visualized the trajectory traces over time, using threshold distances as a color-coded guide (Fig. 5, B-D, Fig. S1-S4). These trace analyses demonstrate that SNX9 PX-BAR sequesters multiple PIP_2_ lipids in its immediate vicinity.

**Fig. 5.**
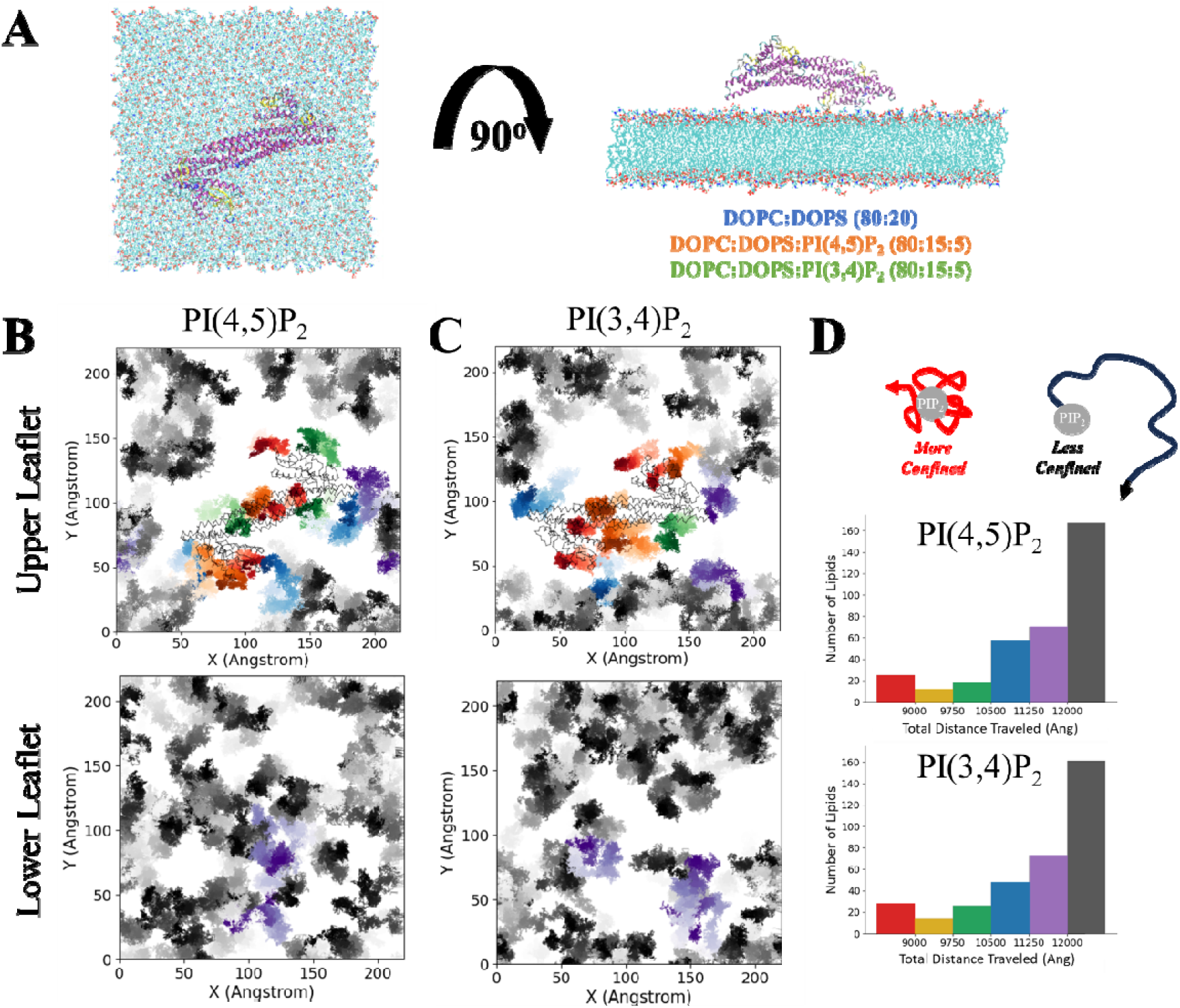
SNX9 PX-BAR locally changes PIP_2_ lipid behavior in atomistic molecular dynamics simulations. **(A)** Top (*Left*) and side (*Right*) views of the initial simulation set up. The membrane is shown in a point style colored by atom type, while the SNX9 PX-BAR domain is represented in a cartoon style colored by secondary structure. Visualization performed with VMD. **(B)** Representative trajectory trace of the last 750 ns (from 1μs trajectory replicate) of the PI(3,4)P_2_ lipids in the upper (*Top*) or lower (*Bottom*) membrane leaflet, colored by total distance traveled. Average position of the SNX9 PX-BAR are shown in wire representation on the upper leaflet. **(C)** Representative trajectory trace of the last 750ns (from 1μs trajectory replicate) of the PI(4,5)P_2_ lipids in the upper (*Top*) or lower (*Bottom*) membrane leaflet, colored by total distance traveled. Average positions of the SNX9 PX- BAR are shown in wire representation on the upper leaflet. **(D)** Histogram of total distances traveled over all replicates for PI(4,5)P_2_ *(Top)* and PI(3,4)P_2_ *(Bottom)*, with bars colored to match trajectory trace colorations in (B) and (C) as well as in Fig. S1-S4.

The sequestration effect of PIP_2_ is most pronounced at the time-averaged positions of the PX domains and the center of the BAR domain (CBD) (Fig. 5, B and C, Fig. S1-S4), with SNX9 PX-BAR overlay). These findings indicate that these regions are most strongly implicated in PIP_2_ binding. While the PX domain has been previously identified as the primary site of PIP_2_ binding, the CBD binding has not been reported in lipid binding. Notably, previous mutational assays have implicated the CBD region in autoinhibition of SNX9, but no lipid selectivity has been reported (*39*). Structural predictions of SNX9 linker region implicated in autoinhibition position the linker region near the CBD. This alignment suggests that the positively charged CBD may mediate nonspecific interactions with PIP_2_ (Fig. S5). Given the plausible non-specific CBD-lipid binding, we focused on the PX domain to investigate the mechanism underlying SNX9’s selectivity for PI(3,4)P_2_ and PI(4,5)P_2_.

We observed that in the upper leaflet of the membranes where SNX9 PX-BAR is bound, there is super-stoichiometric sequestration of PIP_2_ lipids (Fig. 5, B and C). The averaged binding ratios are 3.6 ± 0.5 : 1 for PI(3,4)P_2_ and 3.4 ± 0.8 : 1 for PI(4,5)P_2_ (PIP_2_ : SNX9 PX domain, Table S1).

These binding ratios, exceeding a value of unity, challenge the canonical lock-and-key interaction paradigm that assumes a single PIP_2_ lipid fits into a tight binding pocket. Instead, they suggest that SNX9 PX domain can engage multiple lipids simultaneously through a multivalent binding mechanism. Supporting this, we found that only two out of ten of the individual PX domains in our simulations showed a binding occupancy of less than two lipids (Table S1). The similarity in average binding ratios between PI(3,4)P_2_ and PI(4,5)P_2_ can be attributed to this multivalency, as having multiple binding subdomains that simultaneously bind to the membrane will be the greatest influence on average interactions.

To determine whether the observed superstoichiometric binding is due to specific SNX9 PX-BAR and PIP_2_ interactions rather than the intrinsic properties of the PIP_2_ lipids, we analyzed the average dynamics of the lipids. As shown in Fig. 5, B and C, we observe significant differences in the distance traveled by the PIP_2_ lipids near the SNX9 PX-BAR domain compared to unbound PIP_2_ lipids in the upper leaflet and those in the lower leaflet (Fig. S6). We note that there is no significant difference in the total distance traveled between unbound PIP_2_ lipids in the upper leaflet and in the lower leaflet (Fig. S6). Additionally, no significant spatial variation of PIP_2_ lipids in lower leaflet was observed, which indicates the absence of trans-bilayer coupling due to SNX9 PX-BAR binding. Comparing between PI(3,4)P_2_ and PI(4,5)P_2_ in the lower leaflet, there is no significant difference in the radius of gyration, a measure of lipid motion, indicating that the dynamics of the individual PIP_2_ lipids are similar on average (Fig. S7). Additionally, we quantified the average network connectivity of PIP_2_ clustering as measure of average cluster size and found no significant differences either between the bilayers containing PI(3,4)P_2_ versus PI(4,5)P_2_ nor between the upper (SNX9 bound) and lower (no protein bound) leaflets of each bilayer (Fig. S8). The lack of observable clustering among PIP_2_ is consistent with previous findings, given the ionic composition of our simulations and the local sequestration effect observed (*40*). Taken together, our results indicate that the observed lipid selective sequestration is not driven by intrinsic differences in PI(3,4)P_2_ and PI(4,5)P_2_ behaviors. Instead, the specificity arises from direct interactions between SNX9 PX-BAR and PIP_2_ lipids, particularly those mediated by the PX domain.

To further exclude explanations of the local sequestration based on membrane-mediated effects, we explored possible changes in the behaviors of DOPC and DOPS, as well as changes in overall membrane properties. We found no significant difference in the average radius of gyration and the spatial distributions of DOPC or DOPS in membranes containing or lacking PIP_2_ lipids; put another way, the presence of PIP2 does not locally change the dynamics of other lipids (Fig. S7 and S9). Global membrane properties such as mean curvature and membrane packing defects also showed no significant differences between the three membrane compositions tested in the presence of SNX9 (Fig. S10 and S11). These results indicate that neither the behavior of lipid species present in the membrane nor of the global membrane properties are altered in ways that could explain the observed PI(3,4)P_2_ selectively sequestering by the PX domain of SNX9.

### The Membrane Binding Interface of SNX9 PX domain is Larger Than Previously Identified

We next sought to identify PX-PIP_2_ contacts to determine whether previously characterized canonical binding residues are responsible for PIP_2_ selectivity. By analyzing the frequency of direct contacts between the alpha carbon atoms of residues in the PX domain and the central phosphorus of PI(3,4)P_2_ and PI(4,5)P_2_ (Fig. 6A), we identified three distinct subdomains interacting with PIP_2_. These subdomains are referred to as the insert loop, the canonical pocket, and the fourth helix (Fig. 6 B). We note that the identification of these three subdomains aligns well with the average binding ratios of 3.6 and 3.4 for PI(3,4)P_2_ and PI(4,5)P_2_, respectively, indicating that one or more of these subdomains can interact with multiple PIP_2_ lipids. These subdomains are broad, ranging from 11 to 19 residues with strong membrane interactions, and extend beyond the canonical residues previously identified through sequence alignment (*8*). Between the large membrane interface and observed superstoichiometric binding, the question of selectivity mechanisms should be addressed within each of the subdomains.

**Fig. 6.**
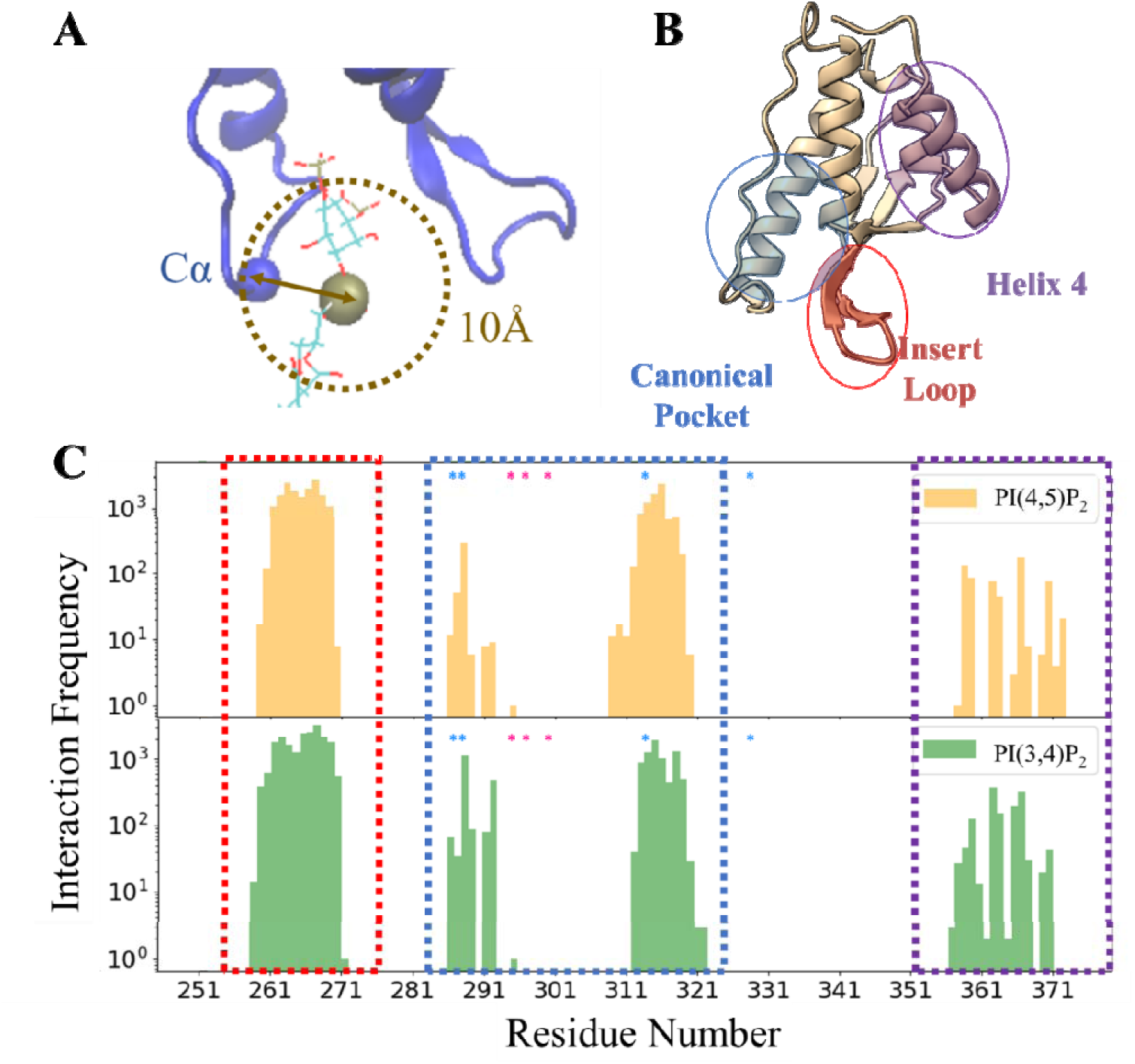
Distinct PIP_2_ binding interfaces of the PX domain can be identified from atomistic molecular dynamics simulations of the SNX9 PX-BAR interacting with membranes containing either PI(3,4)P_2_ or PI(4,5)P_2_. **(A)** Metric for calculating interactions, based on a threshold of 10 Å between the alpha carbon of a residue and the central phosphorus of a PIP_2_ lipid. **(C)** Tabulated log frequencies of interactions for PX domain residues shows clear groupings corresponding to membrane binding areas, designated as the insertion loop (red), the canonical pocket (yellow), and the fourth helix (purple). Blue stars designate canonical binding residues and pink stars designate additional conserved residues identified via homology, according to (*8*). **(B)** Cartoon depiction of the SNX9 PX domain, with approximate membrane binding areas highlighted with ovals; colors correspond to the boxes outlined in (C).

The insert loop subdomain (residues 258-270) is rich in lysines and hydrophobic residues. Its conserved motif, x**K**x(S/T)**K**ΦxGΦ**K**S**YI** (where x represents non-conserved residues, and Φ represents a large hydrophobic residue), is shared by SNX9 and closely related members of the SNX family, SNX18 and SNX33 (Fig. 6C, red dashed box, Fig. S12). The flexibility and composition of this subdomain allows it to penetrate the membrane and interact with negatively charged lipids, anchoring the PX domain to the membrane. It is therefore not surprising that the insert loop shows the highest frequency of contacts with the central phosphate group of both PI(3,4)P_2_ and PI(4,5)P_2_. Unlike the canonical pocket and fourth helix subdomains whose membrane interactions are mediated by specific residues, the PIP_2_ interactions of the insert loop is broadly across its entire length, reflecting its critical role in PIP_2_ interactions (Fig. 6C).

The canonical pocket subdomain (residues 286-293 and 309-320) has been the primary focus in studying PIP_2_ selectivity and binding. Crystal structures of SNX9 PX-BAR domain, along with analogous structural studies of other PX domains, have suggested that a conserved set of residues within the pocket stabilize the PIP headgroup (*8, 11*). However, while the canonical pocket of SNX9 does interface with the PIP_2_ lipids in the membrane, we observed that the conserved residues Tyr294, Arg296, and Lys300 do not interact with the membrane at all (designated with pink * in Fig. 6C). Furthermore, compared to the insert loop, the canonical pocket exhibits greater variation in contact frequency with PIP_2_ among the sequential residues in the loop, due to its more rigid tertiary structure.

The fourth helix subdomain (residues 358-370), which is not traditionally considered as part of the PX domain, lies adjacent to the BAR domain and interacts frequently with the membrane (Fig. 6C, purple dashed box). Like the canonical pocket, this forth helix subdomain exhibits variation in PIP_2_ contact frequency among sequential residues due to its rigid secondary and tertiary structure. While the net charge of the fourth helix is +2, its four positively charged residues (Lys363, Lys367, Arg368, and Arg370) are aligned along its membrane-facing surface. This alignment strongly favor binding to negatively charged lipids such as the PIPs and PS. The alignment of these positively charged residues along the membrane binding surface of the fourth helix suggests that its protruding sidechains could synergistically coordinate the large headgroup of a PIP_2_ lipid.

### The SNX9 PX Fourth Helix Promotes PI(3,4)P_2_ Selectivity

Of the three subdomains identified in this study, only the canonical pocket and the fourth helix interact with PIP_2_ primarily through electrostatic interactions, specifically involving positively charged residues. Because PIP_2_ lipids are strongly negatively charged, - 4 compared to -1 for PI and PS at physiological pH, and have a bulky phosphoinositide headgroup, the specific arrangements of positively charged residues in the canonical pocket and fourth helix are likely key for explaining how the SNX9 PX domain selectively interacts with PI(3,4)P_2_ over PI(4,5)P_2_. The electrostatic map of the membrane binding face of SNX9 PX domain clearly shows the importance of these positively charged residues (Fig. 7A). It also highlights the formation of two distinct pockets of charge surrounding the canonical pocket and the fourth helix.

**Fig. 7.**
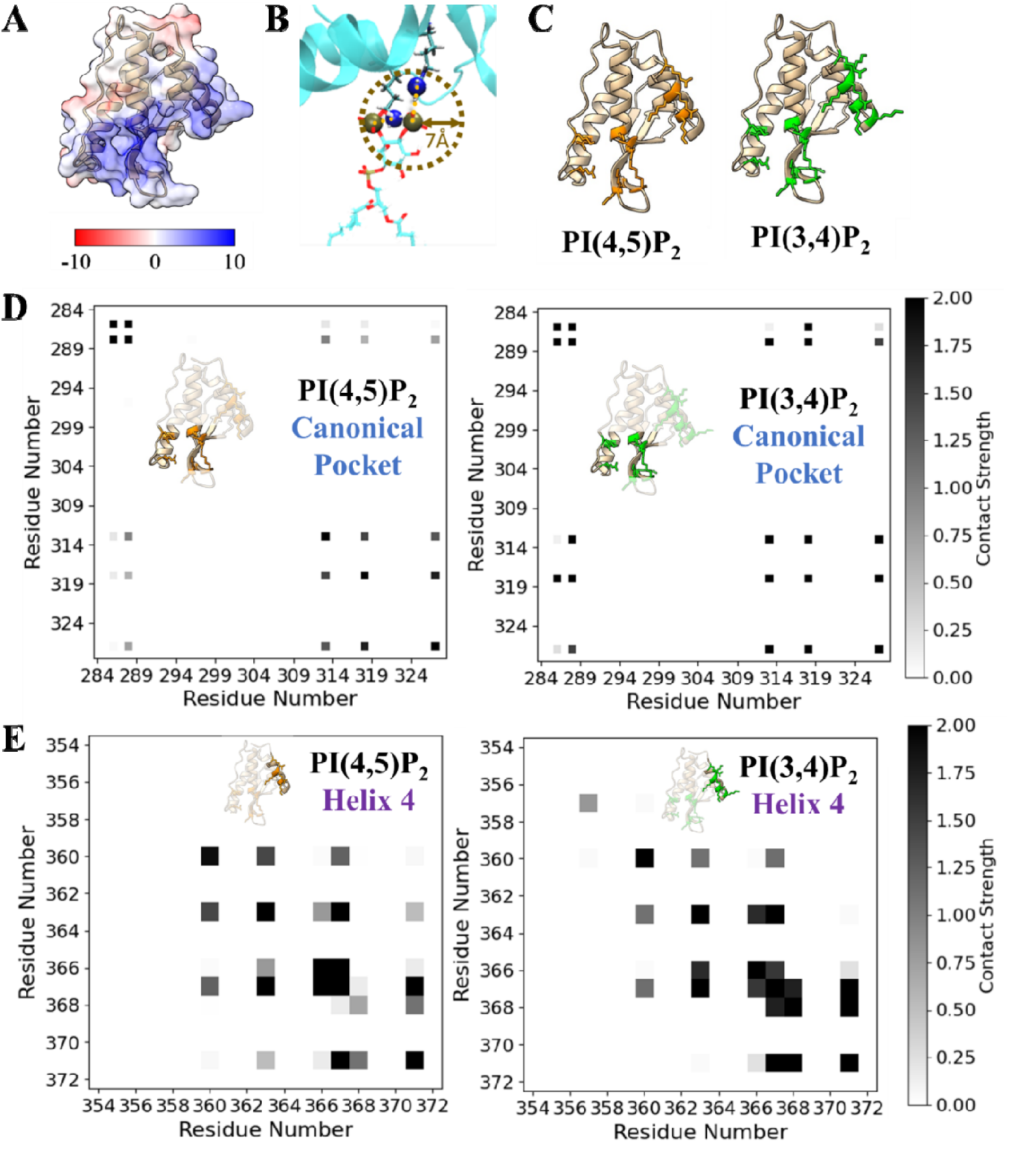
Electrostatics play a primary role in PX-PIP_2_ interactions, particularly in the helix four subdomain. **(A)** Cartoon and electrostatic surface representation of the SNX9 PX domain, with blue indicating positively charged regions and red indicating negatively charged regions. **(B)** Illustration of Bjerrum metric, where a side chain nitrogen atom from an arginine or lysine residue within 7 Å of a PIP_2_ phosphate group was considered as interacting. **(C)** Cartoon representations of the SNX9 PX domain, with super self-coordinating residues for PI(3,4)P_2_ (left) and PI(4,5)P_2_ (right) highlighted in green and orange (respectively) with side chain stick representations. **(D)** PX domain interaction networks for PI(3,4)P_2_ interactions (*Left*) and PI(4,5)P_2_ interactions (*Right*). The subdomains are outlined to highlight the interactions within a subdomain as well as interdomain contacts. **(E)** Fourth helix of the PX domain interaction networks for PI(3,4)P_2_ interactions (*Left*) and PI(4,5)P_2_ interactions (*Right*).

To investigate how these residues could cooperatively bind to the same PIP_2_ lipid, we calculated a contact frequency for all positively charged residues based on the Bjerrum length, considering instances when two residues coordinate the same lipid, though not necessarily the same phosphate group (Fig. 7B, Fig. S13) (*19*). This cooperative frequency based on electrostatic binding groups provides deeper insight into the specificity of lipid interactions. By identifying residues with cooperative self-contact frequencies greater than 1.5, we can determine those involved in major coordination events (Fig. 7C). For the canonical pocket, the major coordinating residues were identical for both PI(3,4)P_2_ and PI(4,5)P_2_, and include Arg286, Lys288, Lys313, and Arg318 (Fig. 7D). However, for the fourth helix, we find a broader set of coordinating residues for PI(3,4)P_2_ over PI(4,5)P_2_. Specifically, residues Lys360, Lys363, Lys366, Arg,367, Lys368 and Arg371 for PI(3,4)P_2_, and Lys363, Lys366, Arg367 and Arg371 for PI(4,5)P_2_, indicating stronger coordinating interactions for PI(3,4)P_2_ (Fig. 7E). Besides, we observe a stronger interaction network between all the residues in the fourth helix for PI(3,4)P_2_ over PI(4,5)P_2_ (Fig. 7E). This reinforces the idea of a more robust coordination for PI(3,4)P_2_, and provides an explanation for how selectivity is maintained, despite superstoichiometric binding ratios. The expansive network of coordinating residues is capable of binding multiple PIP_2_ lipids simultaneously. Furthermore, inter-domain contacts, particularly between the fourth helix and the canonical pocket subdomains, are stronger for PI(3,4)P_2_ than for PI(4,5)P_2_ (Fig. S13). In addition to electrostatic differences, we note that the canonical pocket has a significantly smaller cavity volume of 745 Å^3^ compared to the volume of the fourth helix subdomain of 905 Å^3^ (Fig. S14). Docking predictions further suggest that both PI(3,4)P_2_ and PI(4,5)P_2_ are more likely to bind to the fourth helix subdomain than the canonical pocket (Fig. S14). Collectively, our findings indicate that SNX9 PX domain has a selectivity for PI(3,4)P_2_ over PI(4,5)P_2_, through the fourth helix of the PX domain.

### Specific SNX9-PI(3,4)P_2_ Interactions Protect PI(3,4)P_2_ from Hydrolysis by INPP4B at the Plasma Membrane during Membrane Ruffling

Based on our simulation results showing that SNX9 PX-BAR dimers sequester PI(3,4)P_2_, we hypothesized that SNX9 may influence the hydrolysis of PI(3,4)P_2_ by INPP4B phosphatase. This hydrolysis, converting PI(3,4)P_2_ to PI(3)P, is critical in macropinocytosis for macropinosome closure (*5*). By monitoring PI(3,4)P_2_ enrichment at the plasma membrane of cells overexpressing membrane-bound INPP4B (INPP4B-CAAX), we assessed whether SNX9 protects PI(3,4)P_2_ from INPP4B-mediated hydrolysis through sequestration. Consistent with previous reports, overexpression of wild-type INPP4B (INPP4B-WT), but not the catalytic dead C842S mutant (INPP4B-CS), significantly reduces PI(3,4)P_2_ enrichment at plasma membrane ruffles (Fig. 8, A – C) (*41*). Overexpression of SNX9-mScarlet restores PI(3,4)P_2_ at the plasma membrane of cells expressing INPP4B-WT (Fig. 8, B and D). This result highlights SNX9’s protective role in shielding PI(3,4)P_2_ from hydrolysis by INPP4B.

**Fig. 8.**
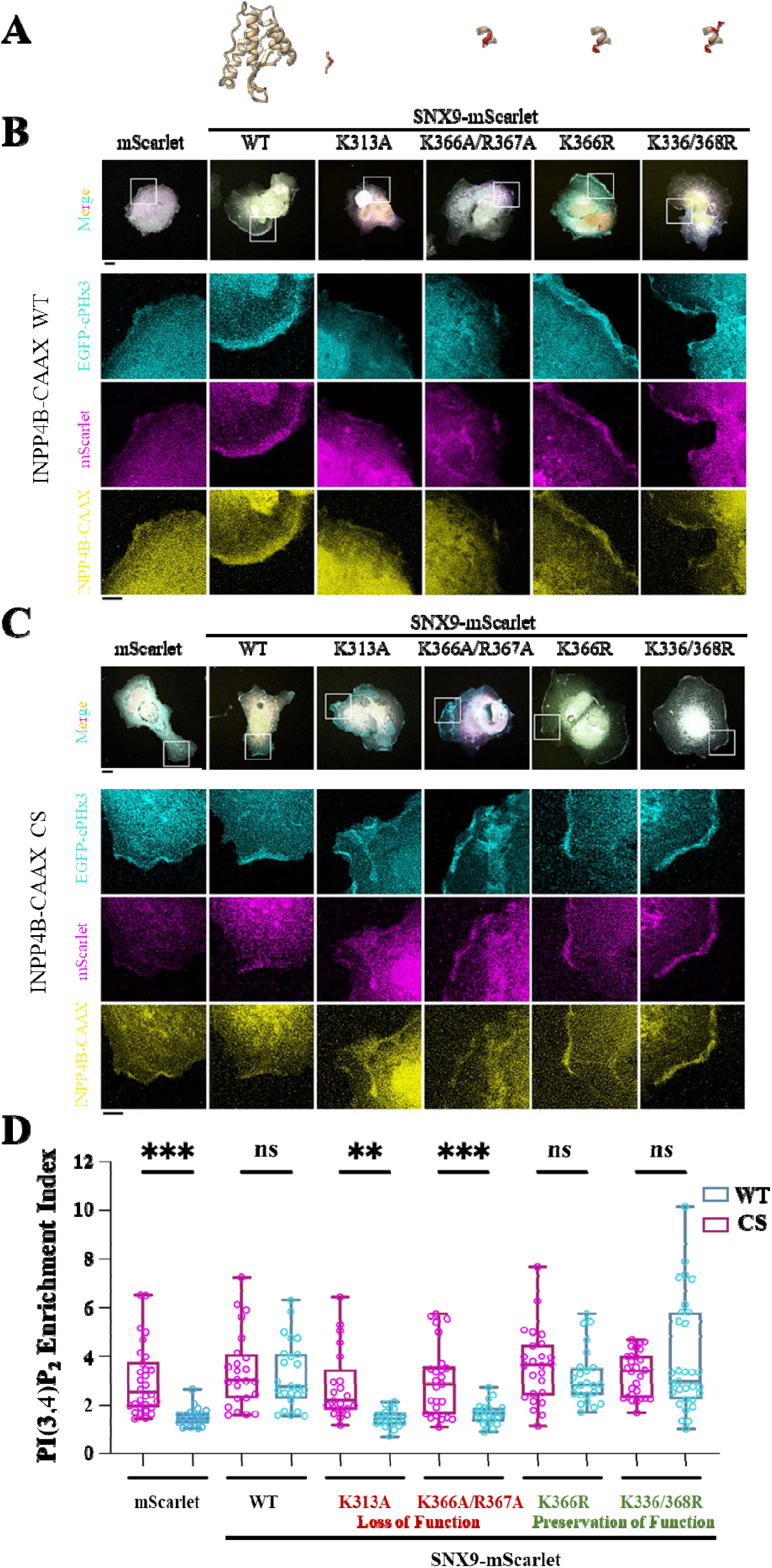
Mutations in the SNX9 PX domain impacts its protective effect on PI(3,4)P_2_ hydrolysis at membrane ruffles. **(A)** Cartoon representations of PX domain, with mutations emphasized by highlighting the affected residue in red and showing the new sidechain. **(B)** Wild-type or **(C)** catalytically dead INPP4B-CAAX, C842S mutant (CS) and PI(3,4)P_2_ reporter were co-transfected into cells together with SNX9-mScarlet or empty vector. After 24 hour incubation in culture medium, cells were fixed and imaged with z-stack confocal microscopy. Projected images were shown. **(D)** PI(3,4)P_2_ enrichment index. The enrichment level of PI(3,4)P_2_ is derived by quantifying the fluorescent intensity fold change at plasma membrane between ruffle and non-ruffle area. Each data point represents one cell. PI(3,4)P_2_ enrichment index in cells expressing indicated SNX9-mScarlet variant together with INPP4B-WT or NPP4B-CS were quantified and compared. Data were analyzed with one-way ANOVA. ***, p < 0.01; ****, p<0.001.

To reveal the functional relevance of SNX9’s PX interaction networks in coordinating PI(3,4)P_2_ that our simulations identified, we performed specific mutations in the SNX9 PX domain, and assessed the consequences of PI(3,4)P_2_ conversion in cellular assays with either functional or non-functional INPP4B phosphatase. We found that loss-of-function mutations in SNX9, K313A in the canonical pocket and K366/367A in the fourth helix, fail to rescue PI(3,4)P_2_ enrichment in cells expressing INPP4B-WT, despite their proper localization at the plasma membrane (Fig. 8, B and D). Conversely, gain-of-function mutations in SNX9, K366R and K366/368R both in the fourth helix, more effectively restored PI(3,4)P_2_ enrichment the plasma membrane. These mutants recapitulated the protective capacity of wild-type SNX9 INPP4B-mediated hydrolysis (Fig. 8D). Notably, in cells expressing the non-functional phosphatase, INPP4B C842S, PI(3,4)P_2_ enrichment index is comparable among all SNX9 variants, indicating that the observed effects of SNX9 on PI(3,4)P_2_ coordination are specific to the functional interaction with INPP4B (Fig. 8D).

## Discussion

We demonstrated that PI(3,4)P_2_ and PI(4,5)P_2_ exhibit distinct spatial distributions upon interacting with SNX9 PX-BAR domain. Although both lipids bind superstoichiometrically to SNX9 via the membrane-remodeling PX-BAR domain, our results revealed that the non-canonical Helix 4 interface in the PX domain is a key determinant of the differential PIP_2_ localization. This finding provides mechanistic insight into how SNX9 can achieve functional lipid selectivity despite its high overall valency for PIP_2_ lipids. Even though the selective binding effect is modest compared to the overall binding of the PX domain, it translates into the preferential spatiotemporal colocalization of SNX9 with PI(3,4)P_2_ rather than PI(4,5)P_2_ at macropinocytic ruffles *in vivo*. Given the generality of the underlying biophysical principles, SNX9’s selective interaction with PIP lipids likely represents a general mechanism for the spatiotemporal recruitment of peripheral membrane proteins in processes like macropinocytosis and endocytosis. As exemplified here by SNX9, many proteins with PIP selective domains, such as PX, PH, FYVE, and C2, might exploit a similar dual mode of PIP interaction. To further elucidate the mechanism behind SNX9’s Helix 4 selectivity for PI(3,4)P_2_ over PI(4,5)P_2_, we characterized a non-canonical binding interface composed of a network of positively charged residues that coordinate the PIP_2_ headgroup. We hypothesized and demonstrated with mutational assays that this Helix 4 interface acts as a molecular checkpoint, preventing the premature hydrolysis of PI(3,4)P_2_ to PI(3)P by INPP4B during macropinocytosis (Fig. 9). Having a checkpoint to prevent the premature conversion of PI(3,4)P_2_ to PI(3)P is especially critical at this juncture in the membrane remodeling process, just prior to membrane fission, to prevent stalled or failed internalization (*32*).

**Fig. 9.**
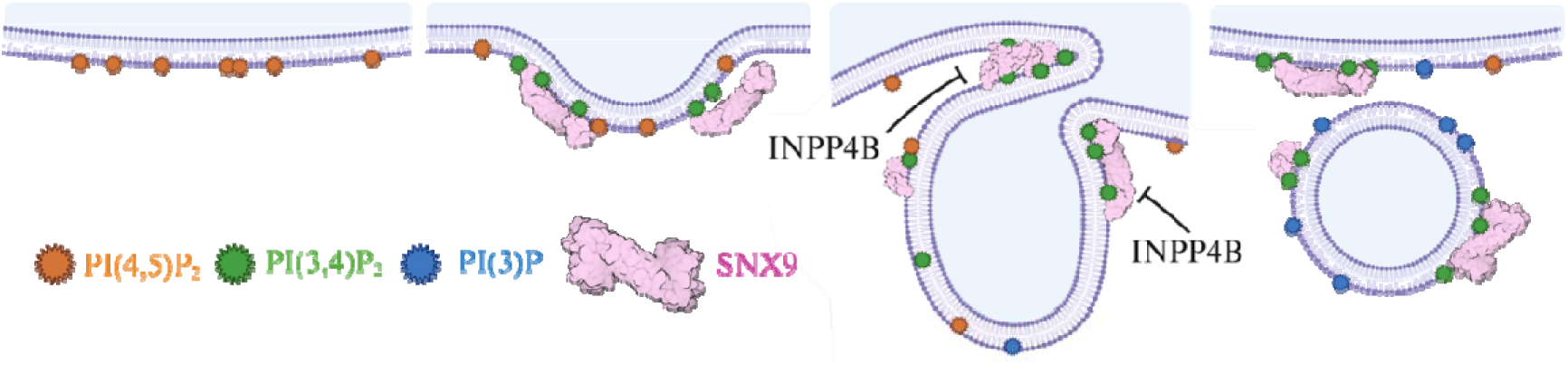
SNX9 prevents the premature hydrolysis of PI(3,4)P_2_ to PI(3)P by INPP4B. As membrane internalization progresses (from *Left* to *Right*), PI(4,5)P_2_ (orange) enrichment at the plasma membrane is followed by the enrichment of PI(3,4)P_2_ (green), while SNX9 (pink) is recruited to the membrane (purple). Once at the membrane, SNX9 colocalizes with PI(3,4)P_2_, shielding it from hydrolysis by INPP4B. After scission of the newly formed vesicle from the plasma membrane, SNX9 dissociates, allowing for the conversion of PI(3,4)P_2_ to PI(3)P to occur.

Using quantitative reconstituted assays, we also showed that full-length SNX9 exhibits membrane curvature sensing and remodeling capabilities, accommodating a wide range of membrane curvatures, independent of the specific PIP_2_ species present. Although SNX9 functions as a relatively weak scaffolding BAR protein, its loosely organized assembly on membrane tubes may allow it to stabilize divers structures – from elongated tubes to constricted necks – which is essential for dynamic processes such as endocytosis and macropinocytosis.

Collectively, our integrated *in silico*, *in vitro* and *in vivo* findings indicate that SNX9’s dual membrane binding domains - its BAR domain for curvature sensing and its PX domain for regulating local PI(3,4)P_2_ concentration – work synergistically for membrane remodeling processes. This dual functionality reinforces SNX9’s central role in membrane dynamics and highlights a broadly applicable mechanism by which peripheral membrane proteins achieve selective recruitment and stabilization at the plasma membrane. Moreover, our Cyro-EM imaging reveals that SNX9-enriched membrane tubes exhibit a broad spectrum of membrane curvatures, with some comparable to those observed in clathrin-mediated endocytosis and dynamin-induced tubes (*30, 42*), suggesting that SNX9 remains membrane-associated throughout the course of remodeling events. Ultimately, SNX9’s ability to coordinate membrane remodeling during lipid conversion with the timely recruitment of downstream effectors via its SH3 domain, such as dynamin and N-WASP, provides a mechanistic basis to ensure coordinated protein recruitment during cellular internalization events. Given the critical role of membrane composition in coordinating membrane remodeling, and considering that enzymatic lipid conversion occurs on a faster timescale than micron-scale membrane remodeling, SNX9’s protective role for PI(3,4)P_2_ against pre-mature hydrolysis to PI(3)P may serve as a temporal checkpoint. This checkpoint ensures that the remodeling process reaches a sufficient level of maturity before proceeding to the final stages of internalization and conversion into an endosomal body. Together, our findings underscore SNX9’s mechanistic role in membrane association and remodeling during lipid conversion, suggesting that SNX9 is both recruited by and orchestrates the maintenance of local PI(3,4)P_2_ concentrations, as well as the timely recruitment of downstream effectors to facilitate membrane remodeling. While not the focus of our study, the previously observed hetero- dimerization of SNX9 with SNX18 could offer a further expanded repertoire of regulatory capabilities (*43*).

The magnitude of the PIP_2_ selectivity of SNX9 PX-BAR domain can be partially attributed to the non-specific superstoichiometric binding of the PX-BAR to the membrane. Binding to multiple lipids simultaneously in non-specific ways can make it difficult to determine selective lipid binding of the PX domain, which may explain why previous *in vitro* literature has reported weak binding for the SNX9 PX domain (*8*). Superstoichiometric binding is likely a general strategy to recruit peripheral proteins to the membrane surface, while specific binding may play a role in the retention of PIP lipids. We note that this does not imply that lipid composition is the exclusive mechanism of recruitment, as other membrane properties such as curvature also change significantly during membrane remodeling events and thus drive protein recruitment. Indeed, the previously reported interplay of the SNX1 PX domain with the BAR domain as partners in membrane recruitment reinforces the idea that curvature plays a likely role in membrane recruitment for other PX-BAR domains, including SNX9 (*44*). Given our results suggesting that SNX9 responds similarly to membranes of similar curvature containing either PI(3,4)P_2_ or PI(4,5)P_2_, we suggest that curvature may be secondary to PIP_2_ selectivity for SNX9 recruitment. In addition, the observed general reduction of lipid mobility around the SNX9 PX-BAR domain due to this superstoichiometric binding may be part of the mechanism of protein-mediated PIP_2_ aggregation, which has been previously reported for other BAR proteins (*45–47*). However, aggregation is a collective effect, and therefore further *in silico* studies should include a much larger membrane system with at least dozens of PX-BAR domains to investigate this phenomenon.

The Helix 4 interface, identified as driving selectivity of SNX9 for PI(3,4)P_2_ over PI(4,5)P_2_, provides the opportunity to reflect more broadly on our treatment of peripheral membrane protein binding modes. We have endeavored to emphasize here that even for lipid selective domains such as the PX domain, multiple lipids are simultaneously bound and so determining selectivity is not always a straightforward task, and requires a detailed look at the structure and dynamics of the domain of interest on a membrane. Sequence data is helpful for identifying conserved residues that may play an important role, but should be paired whenever possible with structural analysis.

The large, negatively charged headgroup of the PIP_2_ lipids define their interactions with peripheral membrane proteins, and distinguishes them particularly from other lipid classes; therefore, the electrostatic interaction mechanism proposed here, coupled with spatial information of the non-canonical binding interface (i.e., how the positively charged sidechains face the membrane and the larger cavity size of the non-canonical interface), is grounded by physical arguments and supported by the mutational assays in cells. While we primarily probed the electrostatic impact by mutating positively charged residues, further work could probe the effect of interface size by introducing bulky, neutral sidechains that would protrude into the adjoining cavity. However, the introduction of such residues could disrupt the protein tertiary structure or lead to insertion of these large neutral residues into the membrane, significantly altering the membrane binding mode as opposed to only probing PIP_2_ selectivity.

It remains unclear what signal eventually drives SNX9 to dissociate from the membrane and free PI(3,4)P_2_ for conversion by INPP4B and other phosphatases. The change in Gaussian membrane curvature as a result of scission could be an important signal to induce a mechanical change in SNX9 organization from a loose organization on tubes to more shallow contacts on spherical vesicles, as has been demonstrated for other BAR domain containing proteins (*48, 49*).

Altogether, it is striking how lipid selectivity of peripheral membrane proteins between two nearly identical lipids, PI(3,4)P_2_ and PI(4,5)P_2_, arises from simple effects such as electrostatics and spatial structural arrangements despite their high valency, and further how selectivity drives the coordination of complex remodeling processes at the plasma membrane. We anticipate that these conclusions are not unique to SNX9 nor the plasma membrane, but rather provide insights into the biophysics behind selective membrane recruitment for a host of peripheral membrane proteins.

## Materials and Methods

### CELL EXPERIMENTS

#### Cell culture, transfection and imaging

COS7 cells were maintained in DMEM supplemented with 10% FBS. Transfection was performed using the plasmid DNA of interest and TransIT (Mirus Bio) according to the manufacturer’s instructions. For live-cell imaging, cells were visualized using a spinning-disc microscope (Carl Zeiss) in imaging medium consisting of phenol red-free DMEM supplemented with 10 mM HEPES and 10% FBS. Medium containing 50 ng/mL PDGF-BB was applied to cells to stimulate membrane ruffles. Images were captured every 2 seconds for 10-15 minutes. For fixed-cell imaging, transfected cells were seeded onto coverslips and fixed with 4% paraformaldehyde, followed by observation with an LSM700 confocal microscope (Carl Zeiss).

#### Image quantification

To analyze the spatiotemporal distribution of PIPs and SNX9, a 1 µm² area was cropped from each membrane ruffle for quantification. The signals were normalized to the highest value and plotted over time. The maximal intensity of SNX9-RFP was set as 100% and aligned to 50 sec for synchronization in figure plotting. For PI(3,4)P_2_ enrichment index quantification, the index is defined as the ratio between PI(3,4)P_2_- enriched membrane and resting plasma membrane. Images were processed with maximal intensity projection, and the contour of PI(3,4)P_2_-enriched membrane regions was manually defined. A similar area of the resting plasma membrane was also defined. The GFP intensity of PI(3,4)P_2_-enriched ruffles and the resting plasma membrane were measured, and the ratio was calculated..

#### Statistical analysis

Statistical analysis was conducted using GraphPad Prism 9.0. Data were analyzed with one-way ANOVA. A P-value of <0.05 was considered statistically significant and is indicated as follows: *, P < 0.05; **, P < 0.01; ***, P < 0.001; ****, P < 0.0001.

### IN VITRO RECONSISTUTION EXPERIMENTS

#### Reagents

Brain L-α-phosphatidylinositol-4,5-bisphosphate (PI(4,5)P_2_, 840046P), 1- heptadecanoyl-2-(5Z,8Z,11Z,14Z-eicosatetraenoyl)-sn-glycero-3-phospho-(1’-myo-inositol-3’,4’- bisphosphate) (17:0-20:4 PI(3,4)P_2_, LM1903) 1,2-distearoyl-sn-glycero-3-phosphoethanolamine- N-[biotinyl(polyethyleneglycol)-2000] (DSPE-PEG(2000)-biotin, 880129P), L-α- phosphatidylcholine (Egg, Chicken) (EPC, 840051), 1,2-di-(9Z-octadécénoyl)-*sn*-glycéro-3- phosphoéthanolamine (18:1 (Δ9-Cis) DOPE, 850725P), 1,2-di-(9Z-octadécènoyl)-*sn*-glycéro-3- phospho-L-sérine (18:1 PS DOPS, 840035), and Cholesterol (700000P), were purchased from Avanti Polar Lipids. BODIPY-TR-C5-ceramide, (BODIPY TR ceramide, D7540), BODIPYFL C5-hexadecanoyl phosphatidylcholine (HPC*, D3803), and Alexa Fluor 488 C5-Maleimide (AX488) were purchased from Invitrogen. Streptavidin-coated polystyrene beads (SVP-30-5) were purchased from Spherotech. β-casein from bovine milk (>98% pure, C6905) and other reagents were purchased from Sigma-Aldrich. Culture-Inserts 2 Well for self-insertion were purchased from ibidi (Silicon open chambers, 80209).

#### Protein purification and labelling

Full length SNX9 construct was a gift from Christian Wunder (Institut Curie, France). The full length SNX9 sequence was cloned into the pProEX HTB plasmid (Invitrogen) between the BamHI and NotI cloning sites, adding a 6xHis tag and rTEV protease cleavage site to the N-terminus of the protein. The plasmid was transformed into Rosetta™ 2(DE3) pLysS (Novagen) and expressed in 2YT medium for 4 hours at 37°C with 0.2 mM IPTG. Cells were lysed by sonication in 50 mM Hepes pH 7.4, 300 mM NaCl, 20 mM Imidazole supplemented with complete EDTA-free protease inhibitor cocktail (Roche). Sample was centrifuged at 20,000 xg and proteins purified using TALON® Metal Affinity Resins (Takara Bio). Protein was eluted with 50 mM Hepes pH 7.4, 300 mM NaCl, 300 mM Imidazole. Proteins were further purified over the Superdex 200 10/300GL column (GE Healthcare) in PBS, 0,5mM EDTA pH 8,0. Pure protein was the labelled with 1:1 protein to dye ratio with Alexa 488 C5 Malemide. Labelled samples were dialyzed into PBS, 0,5mM EDTA pH 8,0, 10% Glycerol, and stored at -80°C.

#### Buffer compositions

The salt buffer inside GUVs, named I-buffer, was 50 mM NaCl, 20 mM sucrose and 20 mM Tris pH 7.5. The salt buffer outside GUVs, named O-buffer, was 60 mM NaCl and 20 mM Tris pH 7.5.

#### GUV preparation

GUVs was prepared using the polyvinyl alcohol (PVA) gel-assisted method (*50*). A PVA solution (5% (w/w) of PVA in a 280 mM sucrose solution) was warmed up to 50°C before spreading on a coverslip, which was cleaned in advance by rinsing with ethanol and MilliQ water. The PVA-coated coverslip was dried in an oven at 60°C for 30 min. 5-10 μl of the lipid mixture (1 mg/mL in chloroform) was spread on the PVA-coated coverslip, followed by drying under vacuum for 30 min at room temperature. The PVA-lipid-coated coverslip was placed in a 10 cm cell culture dish and 0.5 mL of the I-buffer was added on the coverslip, and kept it stable for at least 45 min at room temperature to allow GUV to grow.

#### Sample preparation and observation for measuring dissociation constants

GUVs were incubated with SNX9 at bulk concentrations depending on the designed experiments for at least 15 min at room temperature. Chamber coverslips were passivation with a β−casein solution at a concentration of 5 g.L^-1^ for at least 5 min at room temperature. Experimental chambers were assembled by placing a silicon open chamber on a coverslip. Samples were observed using a Nikon C1 confocal microscope equipped with a X60 water immersion objective.

#### Image analysis

Image analysis was performed by Fiji (*51*). Florescence images were taken at the equatorial planes of GUVs using identical confocal microscopy settings. We measured the fluorescence intensities of SNX9 on spherical GUVs devoid of tubules. The background intensity of the AX488 channel was obtained by drawing a line with a width of 10 pixels perpendicularly across GUV membranes. We obtained the background intensity profile of the line with the x-axis of the profile be the length of the line and the y-axis, the averaged pixel intensity along the width of the line. The background intensity was obtained by calculating the mean value of the sum of the first 10 intensity values and the last 10 intensity values of the background intensity profile. To obtain SNX9 fluorescence intensity on the GUV membrane, we used membrane fluorescence signals to detect the contour of the GUV. A 10 pixel wide band centered on the contour of the GUV was used to obtain the SNX9 intensity profile of the band where the x-axis of the profile is the length of the band and the y-axis, the averaged pixel intensity along the width of the band. SNX9 fluorescence intensity was then obtained by calculating the mean value of the intensity values of the SNX9 intensity profile, following by subtracting the background intensity.

We measured SNX9 surface density on GUV membranes (number of proteins per unit area) by using a previously established procedure (*52*). We related the fluorescence intensity of AX488 to that of a fluorescent lipid (BODIPY FL-C5-HPC, named HPC*). We measure fluorescence intensity of HPC* on GUV membranes at a given HPC* membrane fraction. The surface density of the protein on membranes is 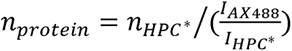, where 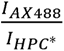 is the factor accounting for the fluorescence intensity difference between HPC* and AX488 at the same bulk concentration under identical image acquisition condition. The area density of HPC*, *ϕ*_HPC*_, can be related to its fluorescence intensity, 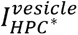, by measuring fluorescence intensities of GUVs composed of DOPC supplemented with different molar ratios of HPC* (0.04-0.16 mole%) and assuming lipid area per lipid is 0.7 nm^2^ (1120 – 4480 HPC* per μm^2^). As such, *n_HPC*_* = *A* × 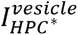, where *A* is a constant depending on the illumination setting in the microscope. We then obtained the surface density of the protein as 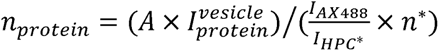, where *n*\* is the degree of labeling for the protein of interest. Finally, we obtained the surface fraction of the protein *Ф_protein_* = *n_protein_* x *a_protein_*, where *a_protein_* is the area of a single protein on membranes, *a_BAR domain_* ≌ 50 nm^2^ (*34*).

To obtain the dissociation constant of SNX9 on GUV membranes, we fitted the surface densities of SNX9 *ϕ_v_*, determined from the fluorescence signals, as a function of SNX9 bulk concentration *C̅_bulk_* to

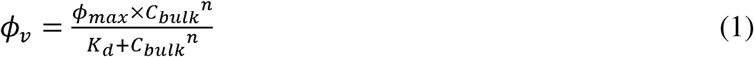

where *ϕ_max_* is the maximum surface density, *K_d_* is the dissociation constant and *n* is the Hill coefficient.

##### Tube pulling experiments

The experiments were performed on a setup with a Nikon C1 confocal microscope equipped with a X60 water immersion objective, micromanipulators for positioning micropipettes and optical tweezers as previously described (*52*). To pull a tube, a GUV was held by a micropipette, brought into contact with a streptavidin-coated bead trapped by the optical tweezers, followed by moving away from the bead. Tube radius *R* was measured by the ratio of lipid fluorescence intensity on the tube and on the GUV as 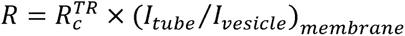, where 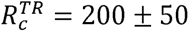 nm is the previously obtained calibration factor for using BODIPY TR ceramide as lipid fluorescence reporter in the same setup by performing a linear fit of membrane fluorescence ratio (*I_tube_*/*I_vesicle_*)_membrane_ and lipid radii *R* measured by *R* = *f*/(4*πσ*), where 1 is the force applied by the optical tweezers to hold the tubes and *f* the membrane tension tuned by the micropipette holding the GUVs (*34, 52*).

To obtain protein/membrane fluorescence intensity in tube pulling experiments, we defined a rectangular region of interest (ROI) around GUV membranes and around the membrane tube such that the membrane/tube were horizontally located at the center of the ROI. We then obtained an intensity profile along the vertical direction of the ROI by calculating the mean fluorescence intensity of each horizontal line of the rectangle. To account for protein fluorescence outside the GUVs, the background protein intensity was obtained by calculating the average value of the mean of the first 15 intensity values from the top and from the bottom of ROI. A similar procedure was used the membrane background. The protein/membrane fluorescence intensities were obtained by subtracting the background intensity from the maximum intensity value in the intensity profile.

##### SNX9 sorting data analysis

We fit the sorting data using the curvature mismatch model (*35*). The free energy of the system is

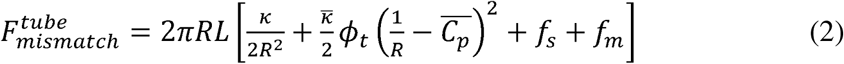

Where *R* is the tube radius, *L* is the tube length, *κ* is the bending modulus of the protein-bound membrane, *κ* is the bending modulus of the protein-free membrane, *κ* is an elastic coefficient penalizing mismatch between protein and membrane curvature, *ϕ_t_* is the protein areal fraction on the tube, 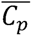 is a phenomenological coefficient related to the membrane-bound protein’s intrinsic curvature, and *f_s_* and *f_m_* are the energy densities of membrane stretching and protein mixing entropy on membranes. At equilibrium the chemical potentials of the lipids and proteins on the GUV and on the tube are equal, thus an implicit dependence of *ϕ_v_* (protein area fraction on the GUV) on the tube curvature can be written as

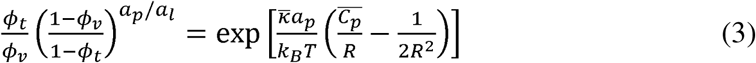

where *a_p_* and *a_l_* is the membrane areas occupied by a membrane bound protein and a lipid, respectively. *S* = *ϕ_t_*/*ϕ_v_* is the sorting ratio and *C* = 1/*R* the tube curvature. By fitting the sorting data with this equation, one can obtain *κ* and 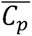.

##### Cryo-electron microscopy experiments

A heterogeneous solution of LUVs was generated by resuspension in buffer (60 mM NaCl and 20 mM Tris pH 7.5) of a dried lipid film composed of 80 % DOPC, 10 % DOPS and either 10 % PI(4,5)P2 or 10 % PI(3,4)P2 (molar ratios). 1 µM of SNX9 was incubated with the lipid suspensions at 0.1 mg.mL^-1^ at room temperature for an hour. The samples were vitrified on copper holey lacey grids (Ted Pella) using an automated device (EMGP, Leica) by blotting the excess sample on the opposite side from the droplet of sample for 4 s in a humid environment (90% humidity). Imaging was performed by a Glacios microscope (Thermofisher) running at 200 kV and equipped with a Falcon IVi direct detector (Thermofisher).

### ATOMISTIC MOLECULAR DYNAMICS

The initial structure of the truncated SNX9 PX-BAR dimer (residues 201-595) was generated with AlphaFold2 (*38*) to model missing loops and the N-terminal amphipathic helix. The predicted structure was compared to previously solved structures (PDB IDs: 2RAI, 2RAK), and found to have only small deviations from each structure (Figure 1) (*11*). The PX-BAR dimer was oriented parallel to the membrane and translated to initially be 3nm above the membrane surface. Three sets of molecular dynamics simulations were conducted on membranes that were initially 22 nm by 22 nm in the xy plane, with compositions of 80:20 mole percent DOPC:DOPS, 80:15:5 mole percent DOPC:DOPS:PI(4,5)P_2_, and 80:15:5 mole percent DOPC:DOPS:PI(3,4)P_2_, and solvated with 0.15M KCl; five replicates of each membrane composition were conducted. Atomistic systems were generated with the CHARMM-GUI platform (*53, 54*), and simulated with the CHARMM-36m force field using GROMACS 2021.5 (*55*) for 1000ns per replicate in the constant NPT ensemble after standard equilibration, for a total of 5 μs per system. Analysis was conducted over the last 750ns of each replicate in Python with MDAnalysis (*56*). Visualization was conducted with ChimeraX (*57*) and Visual Molecular Dynamics (*58*).

## Acknowledgments

We thank Olena Pylypenko and Korbinian Liebl for insightful discussion and Christian Wunder for SNX9 plasmid. F.-C.T. and P.B. is a member of the CNRS consortium Approches Quantitatives du Vivant (AQV), Labex Cell(n)Scale (<u>ANR-11-889 LABX0038</u>) and Paris Sciences et Lettres (<u>ANR-10-IDEX-0001-02</u>). The authors greatly acknowledge the Cell and Tissue Imaging (PICT-IBiSA), Institut Curie, member of the national infrastructure France- BioImaging (https://ror.org/01y7vt929) supported by the French National Research Agency (ANR-24-INBS-0005 FBI BIOGEN). PB team is supported by the Fondation pour la Recherche Médicale (FRM) (FRM EQU202003010307) and by the European Union (ERC, PushingCell, #101071793). Views and opinions expressed are however those of the authors only and do not necessarily reflect those of the European Union or the European Research Council. Neither the European Union nor the granting authority can be held responsible for them. Simulations were performed using computing resources provided by the University of Chicago Research Computing Center (RCC), the Department of Defense High Performance Computing cluster (HPCMP), and the National Science Foundation ACCESS cluster.

## Funding

Labex Cell(n)Scale (F-CT)

Université Paris Sciences et Lettres-QLife Institute ANR-17-CONV-0005 Q-LIFE (MyoMemActin) (F-CT)

Agence Nationale pour la Recherche (ANR-20-CE11-0010-01, ActinFission) (F-CT) Institut Curie and the Centre National de la Recherche Scientifique (CNRS) (F-CT and PB) Fondation pour la Recherche Médicale (FRM) (FRM EQU202003010307) (F-CT and PB) ERC (PushingCell, #101071793) (F-CT and PB)

National Science and Technology Council (Republic of China, Taiwan) (Y-WL)

National Institute of General Medical Science (NIGMS) of the National Institutes of Health (NIH) R01GM063796 (GAV and JRB)

University of Chicago France And Chicago Collaborating in The Sciences (GAV and JRB) Chateaubriand Science and Technology Fellowship (JRB)

## Author contributions

JRB, Y-WL, GAV, and F-CT designed the initial project.

JRB, CJS, SSL, NdV, AB and F-CT performed experiments and analyzed results with feedback from Y-WL, GAV and PB.

JM and SA purified proteins.

JRB and F-CT wrote the original draft with inputs and revisions from AB, PB, Y-WL and GAV.

Conceptualization: JRB, Y-WL, GAV, and F-CT

Methodology: JRB and F-CT

Investigation: JRB, CJS, SSL, NdV, AB, F-CT

Resources: JM, SA

Visualization: JRB, F-CT

Supervision: Y-WL, GAV, F-CT

Writing—original draft: JRB, F-CT

Writing—review & editing: JRB, F-CT, AB, JM, SA, Y-WL, GAV, PB

Funding acquisition: FCT, Y-WL, GAV, PB

## Data and materials availability

All data needed to evaluate the conclusions in the paper are available in the main text and the Supplementary Materials.

## Supplementary Materials

See corresponding document.

