## SupplementalTablesandFigures for "A Non-Canonical Interface of SNX9 PX Domain Selectively Sequesters PI(3,4)P_2_ Lipids, Protecting Them from Hydrolysis"

Jeriann R. Beiter *et al.*

**This PDF file includes:**

Table S1
Figs. S1 to S14

| | PI(3,4)$P_2$ | | | | |
|---|---|---|---|---|---|
| | **BAR** | **PX1** | **PX2** | ***Total*** | ***PX avg*** |
| **Replicate 1** | 3.53 | 4.17 | 3.56 | 11.26 | 3.87 |
| **Replicate 2** | 4.76 | 3.24 | 2.93 | 10.93 | 3.09 |
| **Replicate 3** | 5.98 | 3.56 | 3.99 | 13.53 | 3.78 |
| **Replicate 4** | 5.66 | 2.00 | 4.15 | 11.81 | 3.08 |
| **Replicate 5** | 3.75 | 4.31 | 0.63 | 8.69 | 2.47 |
| | **PI(4,5)$P_2$** | | | | |
| | **BAR** | **PX1** | **PX2** | ***Total*** | ***PX avg*** |
| **Replicate 1** | 5.31 | 2.55 | 4.60 | 12.46 | 3.56 |
| **Replicate 2** | 3.11 | 3.25 | 2.79 | 9.15 | 3.02 |
| **Replicate 3** | 6.08 | 4.22 | 3.28 | 13.58 | 3.75 |
| **Replicate 4** | 2.43 | 4.09 | 4.90 | 11.42 | 4.50 |
| **Replicate 5** | 3.06 | 2.46 | 1.08 | 6.60 | 1.77 |

**Table S1.** $PIP_2$ Occupancy values for the SNX9 PX-BAR dimer.

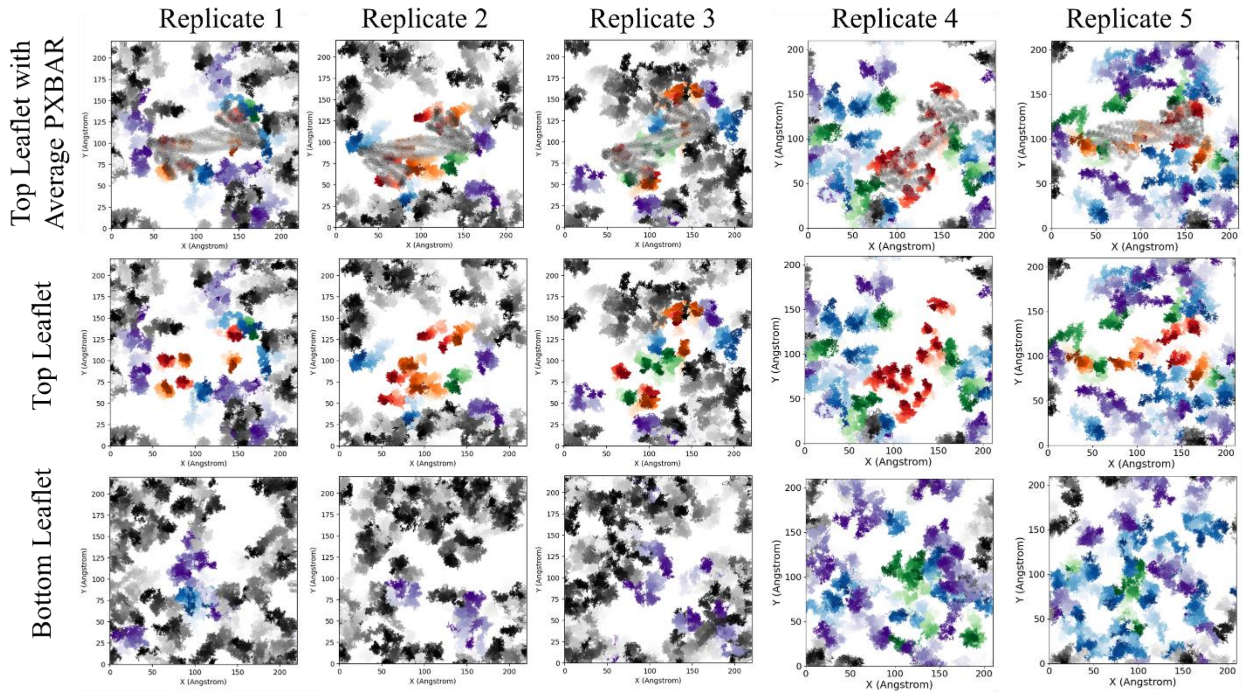


**Fig. S1.** PI(3,4)$P_2$ trajectory trace plots for all replicates.

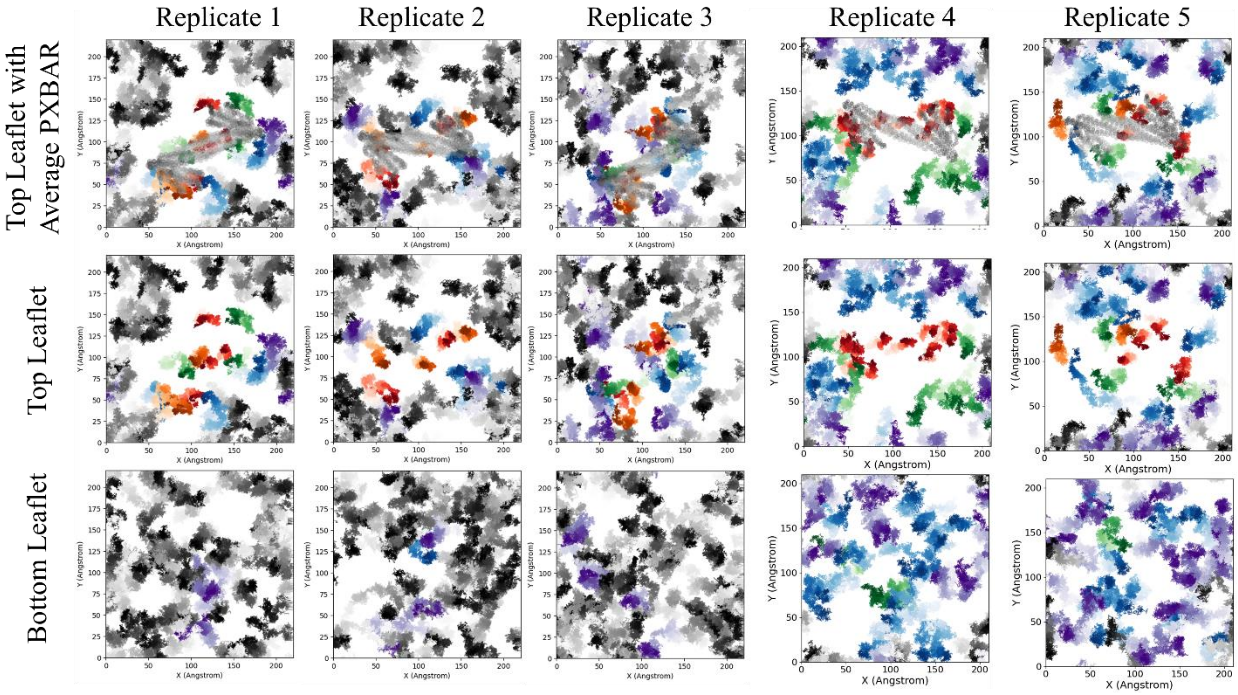


**Fig. S2.** PI(4,5)$P_2$ trajectory trace plots for all replicates.

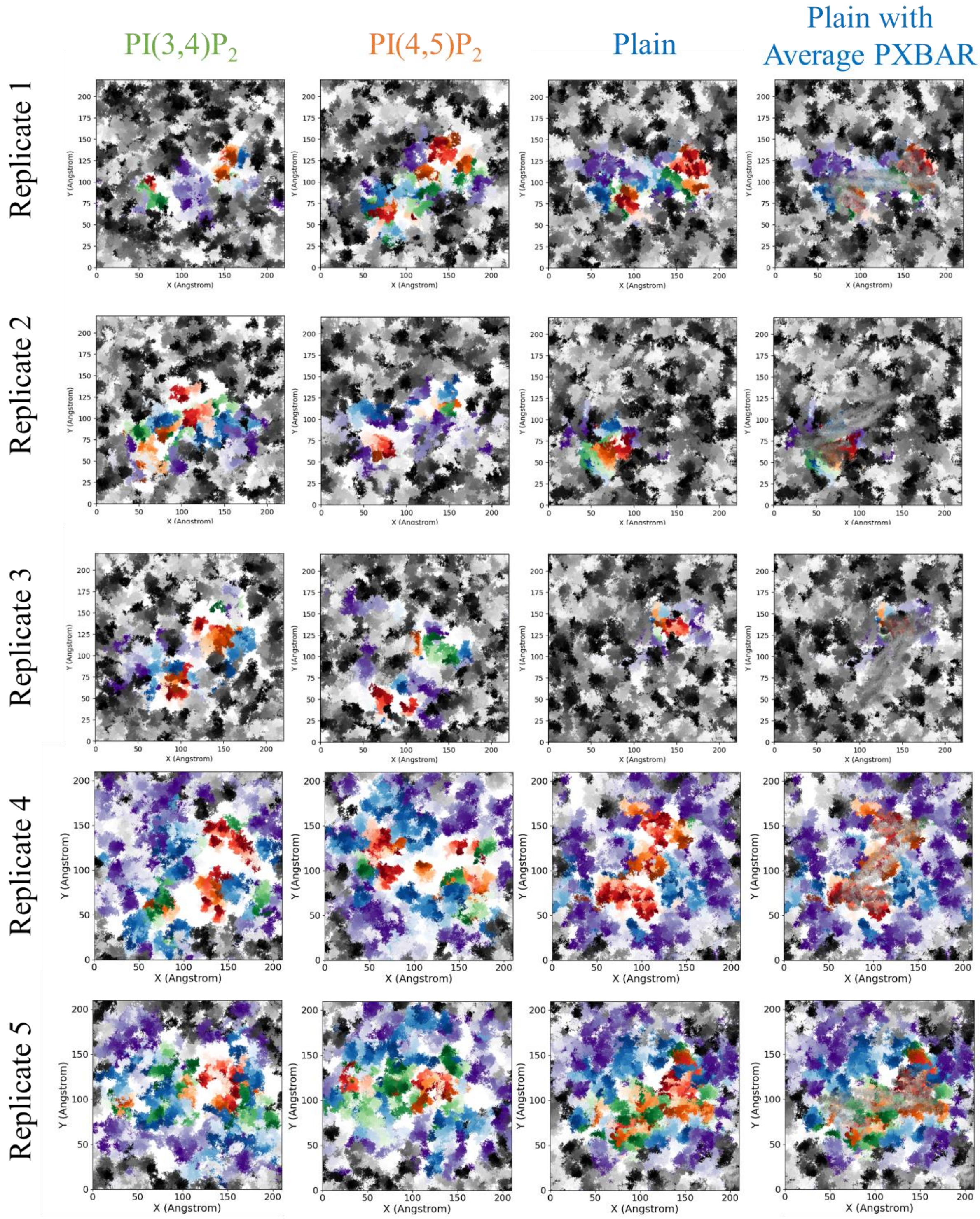


**Fig. S3.** DOPS trajectory trace plot for the top leaflet in all replicates.

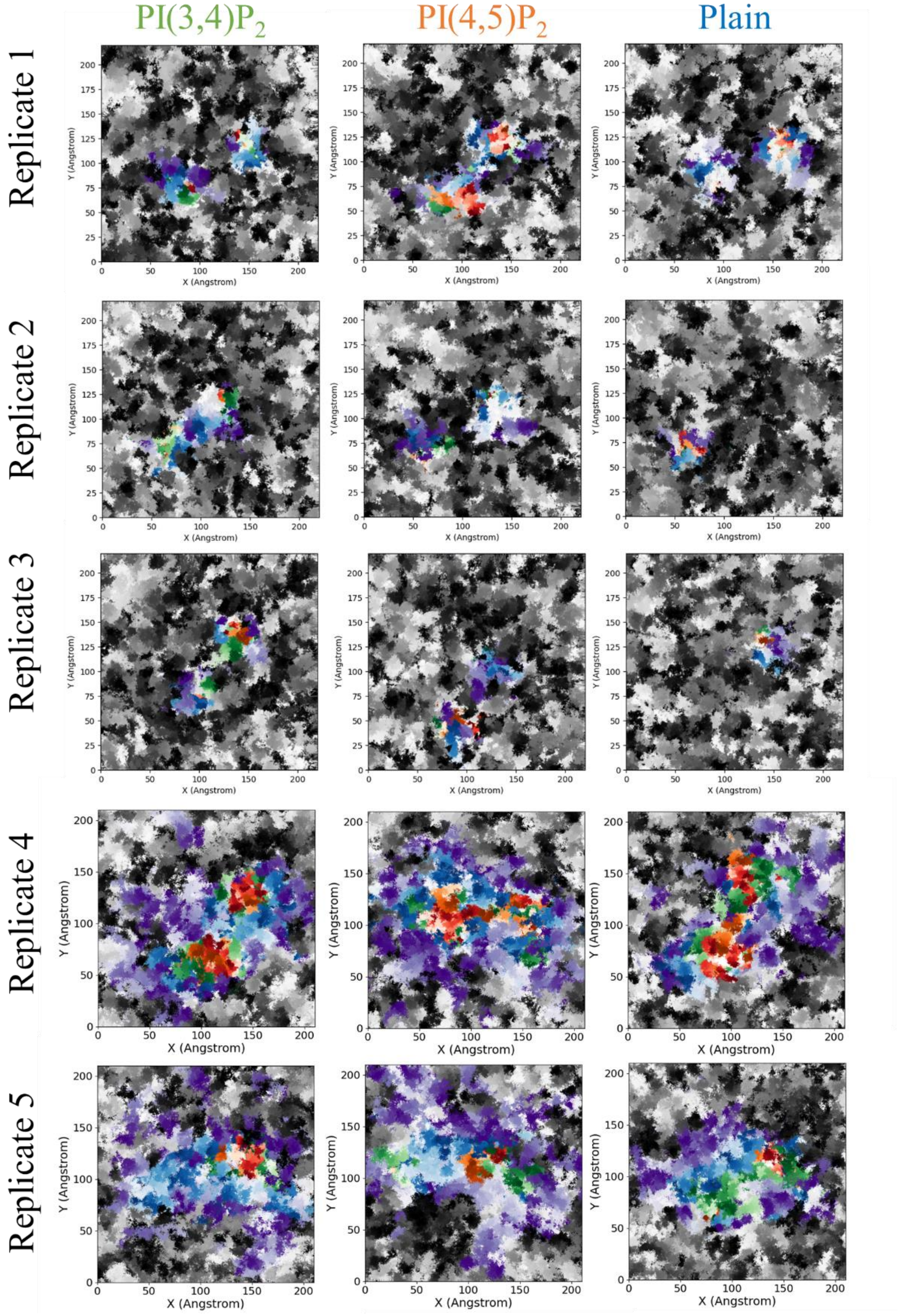


:q

**Fig. S4.** DOPC trajectory trace plot for the top leaflet in all replicates.

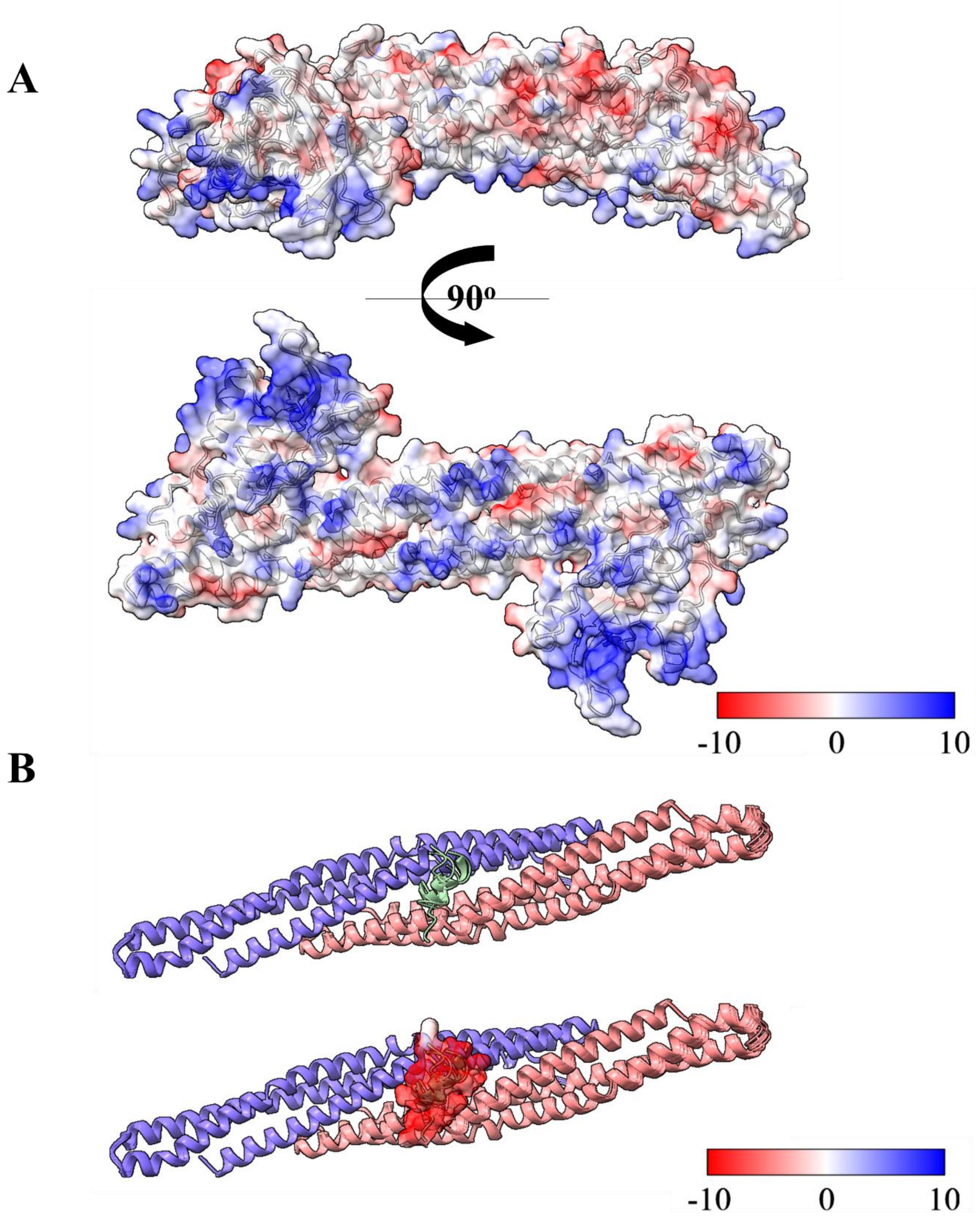


**Fig. S5.** Electrostatic Map of the PXBAR domain and autoinhibition motif. **(A)** Electrostatic surface map shown transparently over a cartoon representation of the SNX9-PXBAR domain from the side (top) and membrane binding interface (bottom). **(B)** Cartoon (top) and surface (bottom) depiction of predicted BAR domain (blue/pink) and autoinhibition motif (green) interaction from AlphaFold2. Visualization generated with ChimeraX.

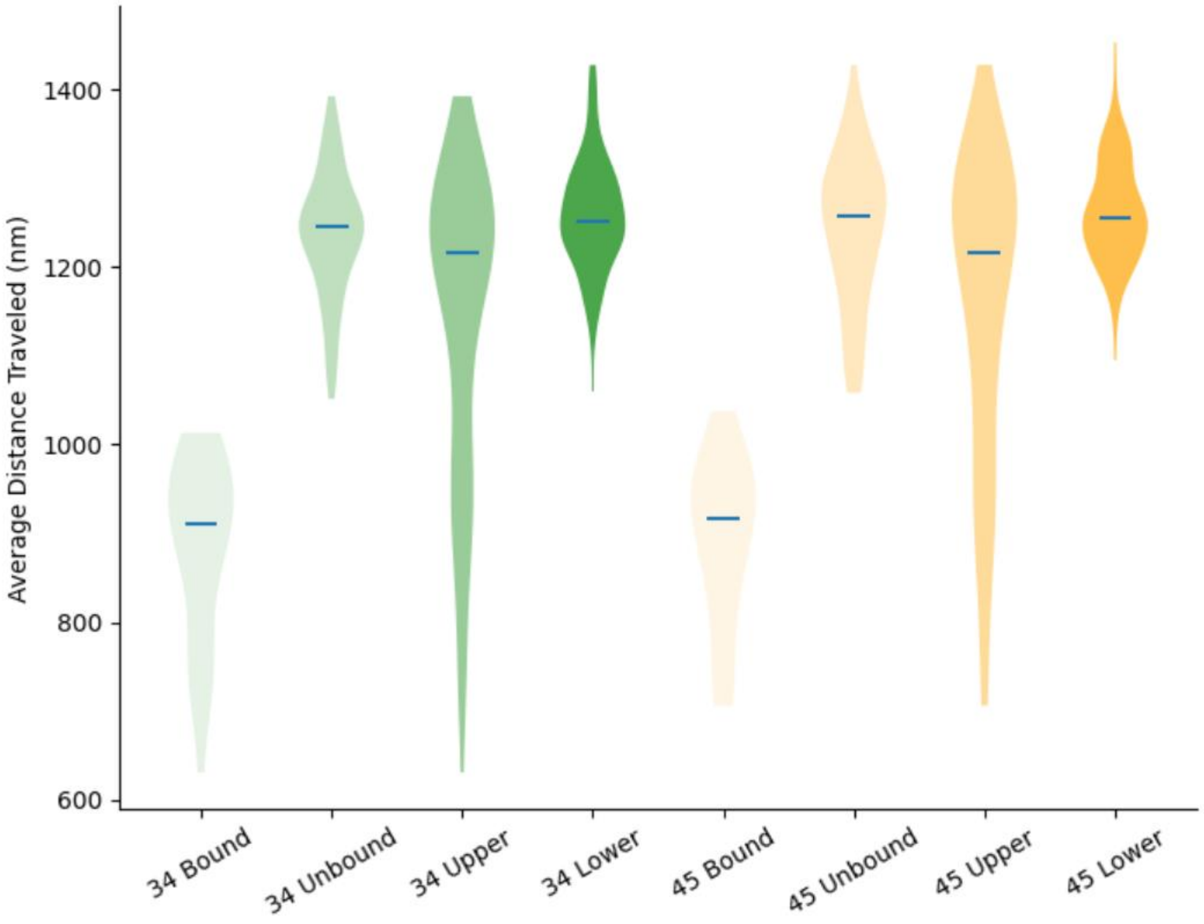


**Fig. S6.** Composite violin plot of average distance traveled of $PIP_2$ lipids.

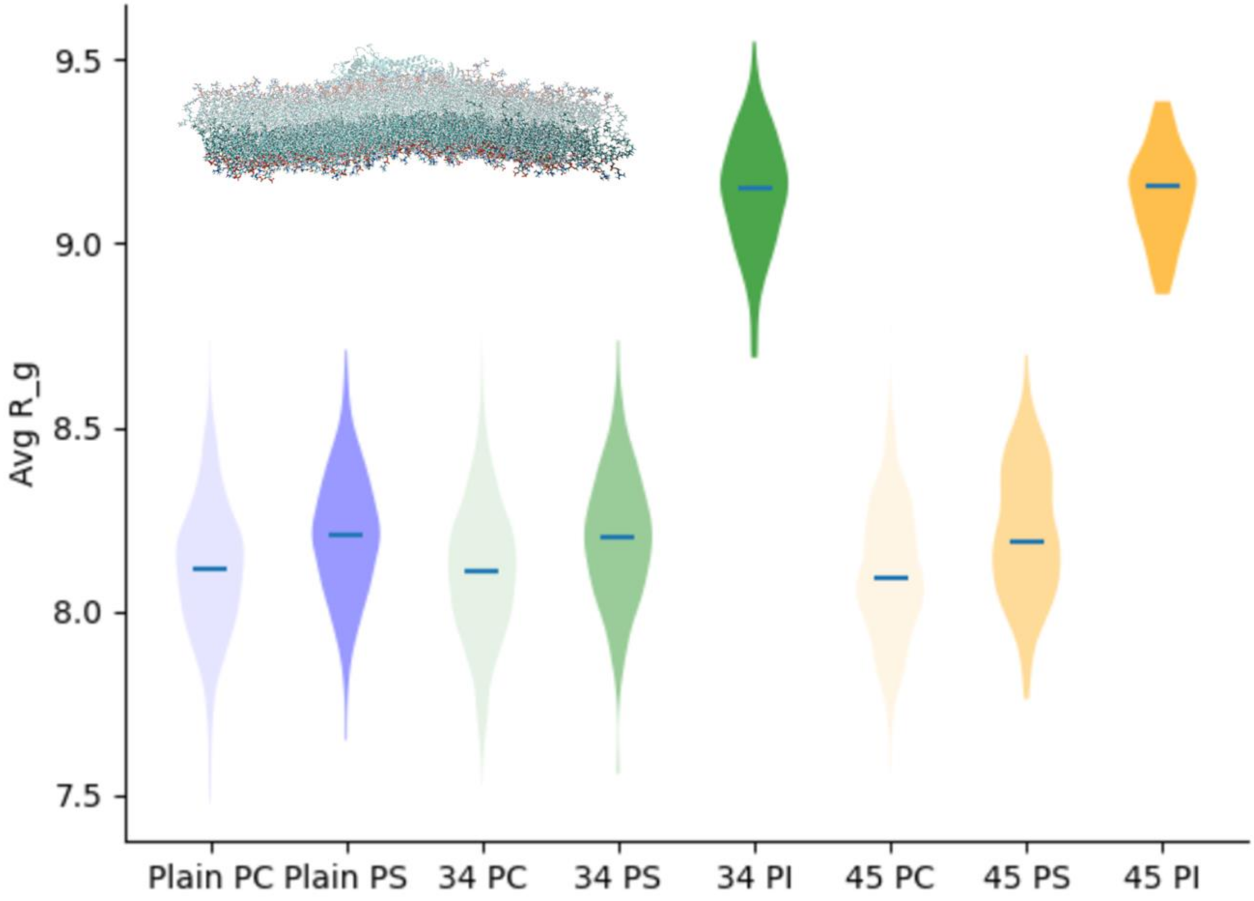


**Fig. S7.** Average Radius of Gyration of lipids in bottom leaflets across replicates.

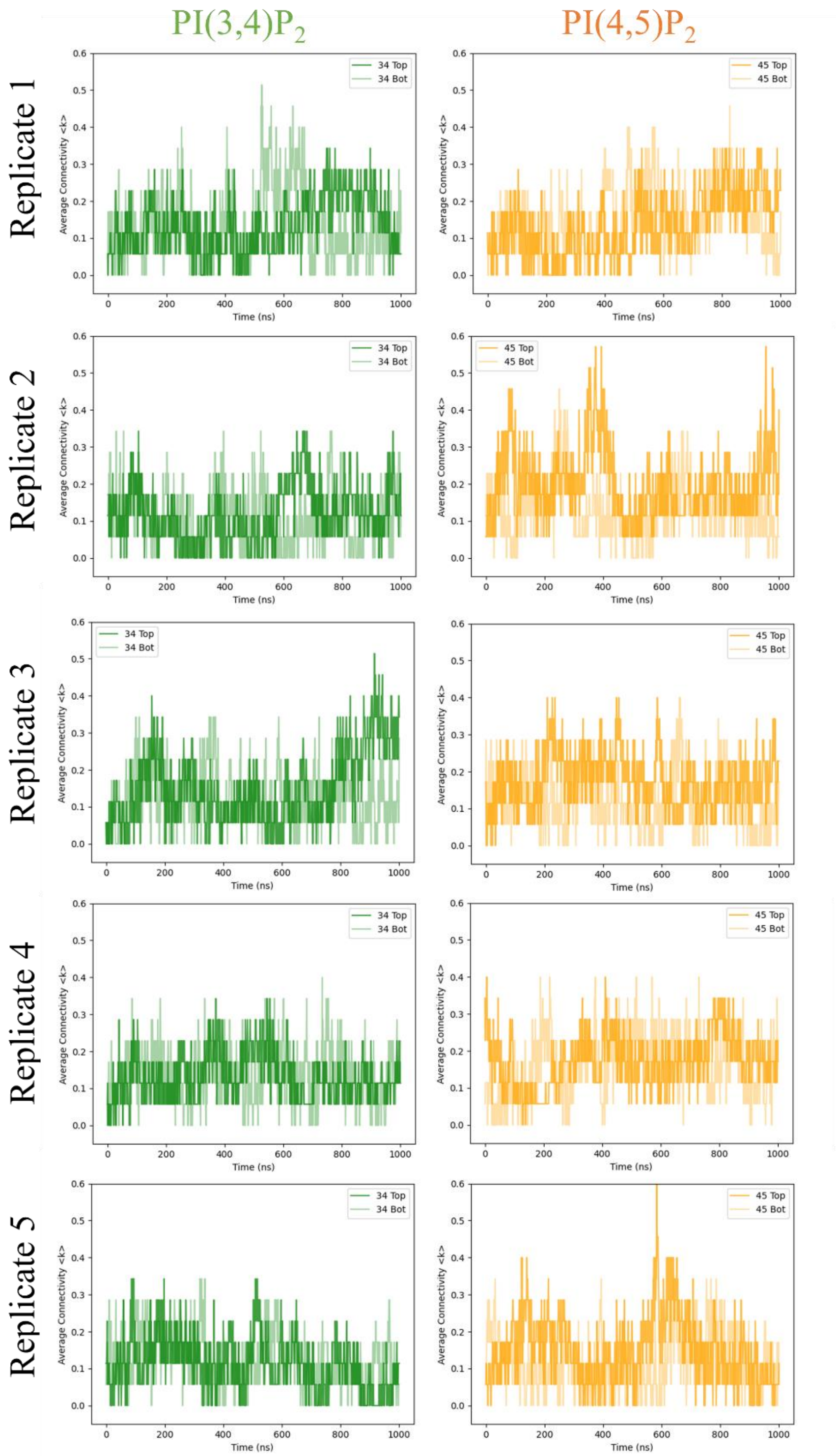


**Fig. S8.** Average connectivity <k> values for PI(3,4)$P_2$ and PI(4,5)$P_2$ as a function of time over the course of each replicate. Dark green or orange indicates the clustering values for top leaflet, while light green or orange indicates the clustering values for the bottom leaflet.

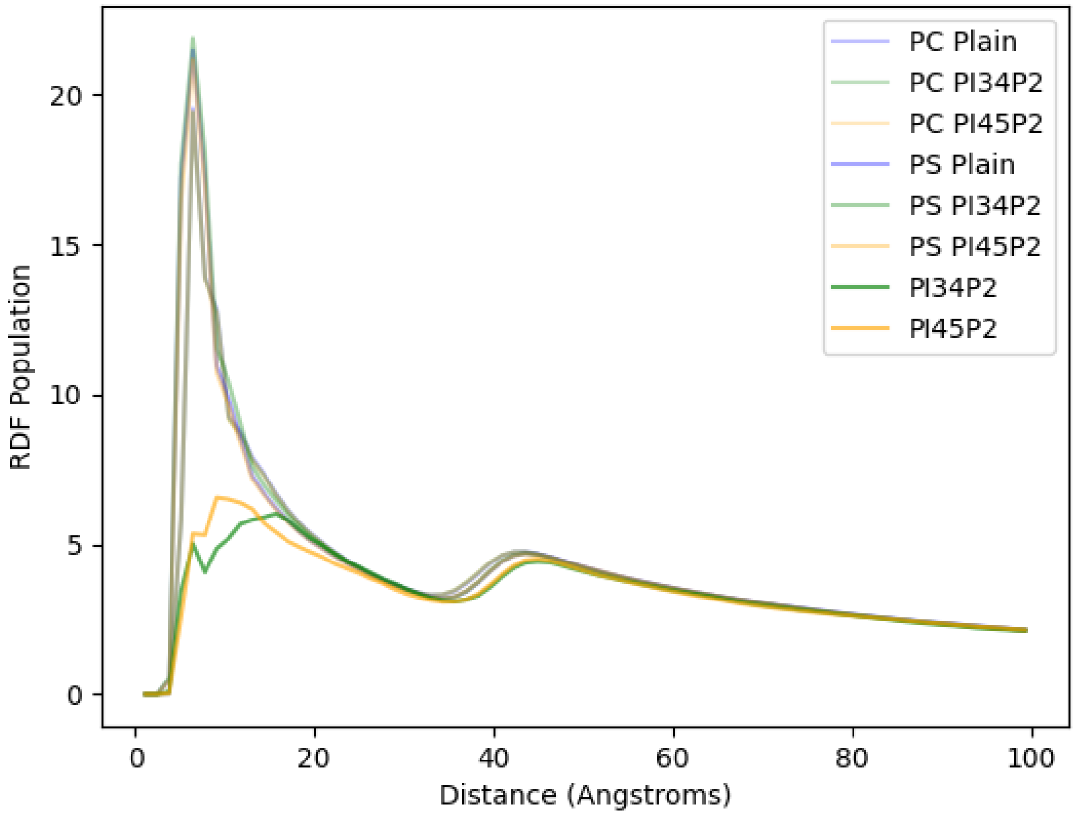


**Fig. S9.** Composite radial distribution function for all lipids.

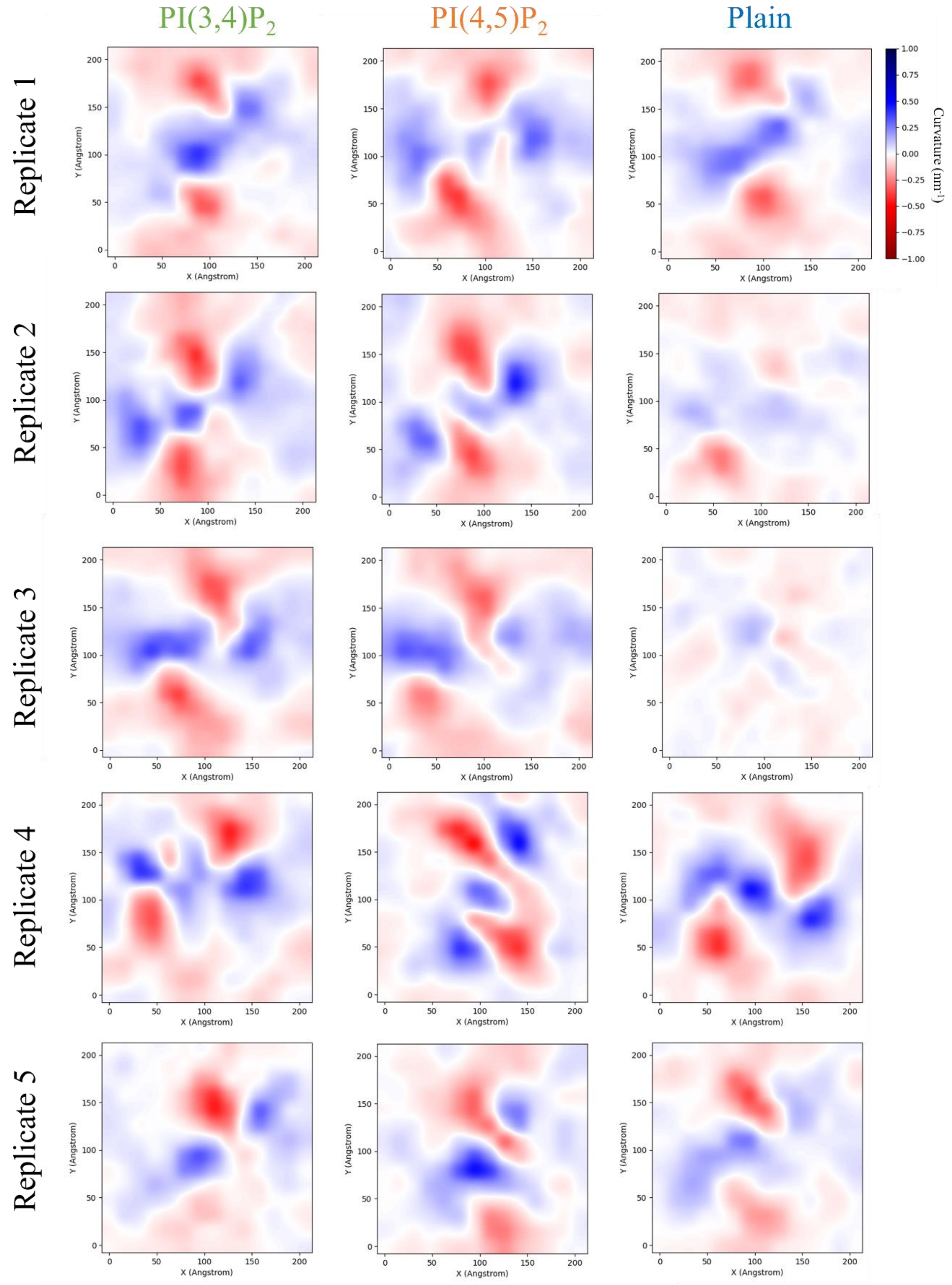


**Fig. S10.** Curvature analysis for both the upper leaflet of each replicate.

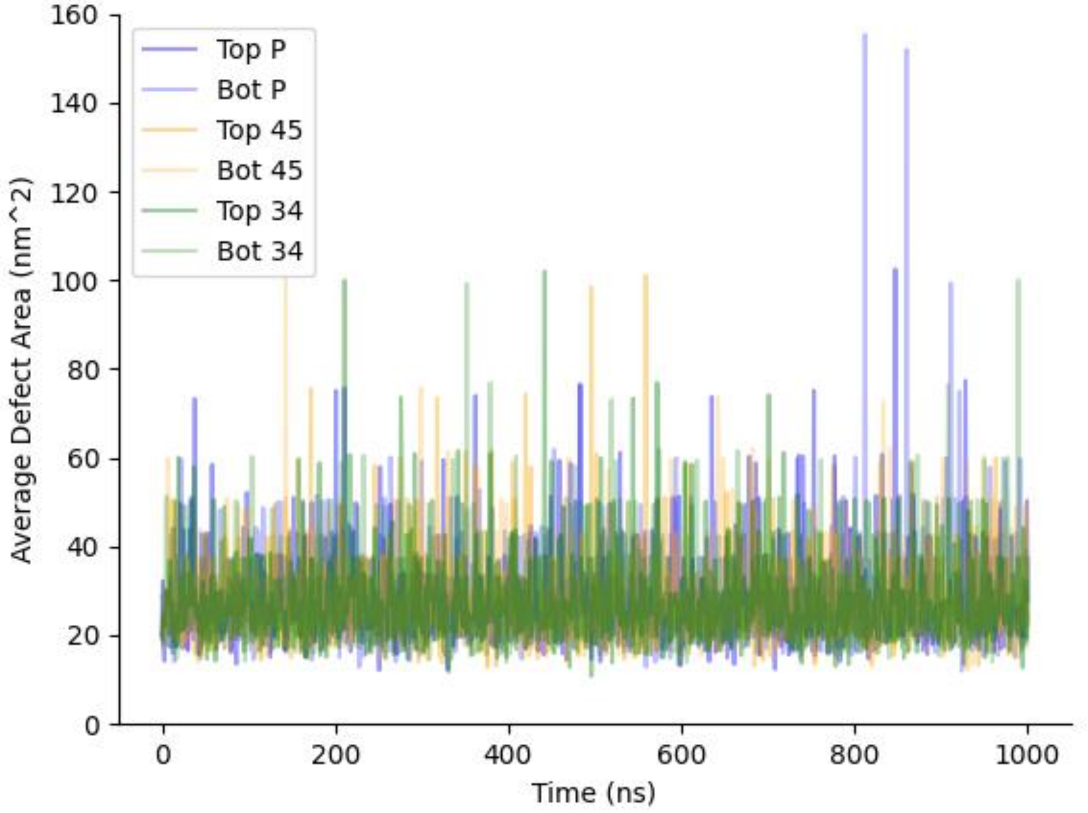


**Fig. S11.** Average membrane defect area for both the upper and lower membrane leaflets.

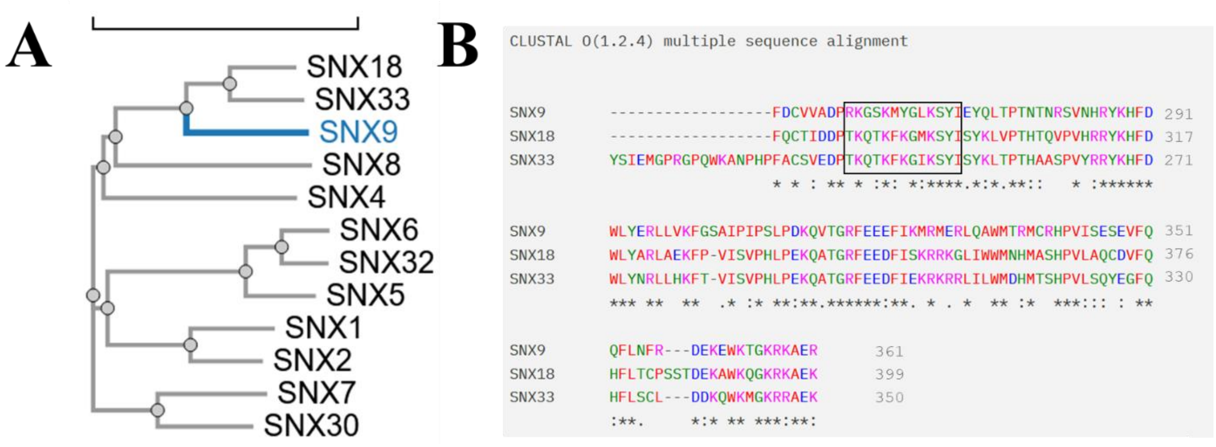


**Fig. S12.** Phylogeny analysis and sequence alignment of PX domains of closely related PXBAR containing SNX proteins, conducted using the CLUSTAL OMEGA MSA program (59). **(A)** Phylogeny of PX domains from all human SNX-PXBAR proteins, demonstrating highest sequence conservation between SNX9 to SNX18 and SNX33. **(B)** MSA of PX domains of SNX9, SNX18, and SNX33, with color to indicate amino acid properties such as charge or hydrophobicity. * indicates an amino acid identically conserved in all three proteins; : indicates a higher conservation of property but not identity; and . indicates a lower conservation of property.

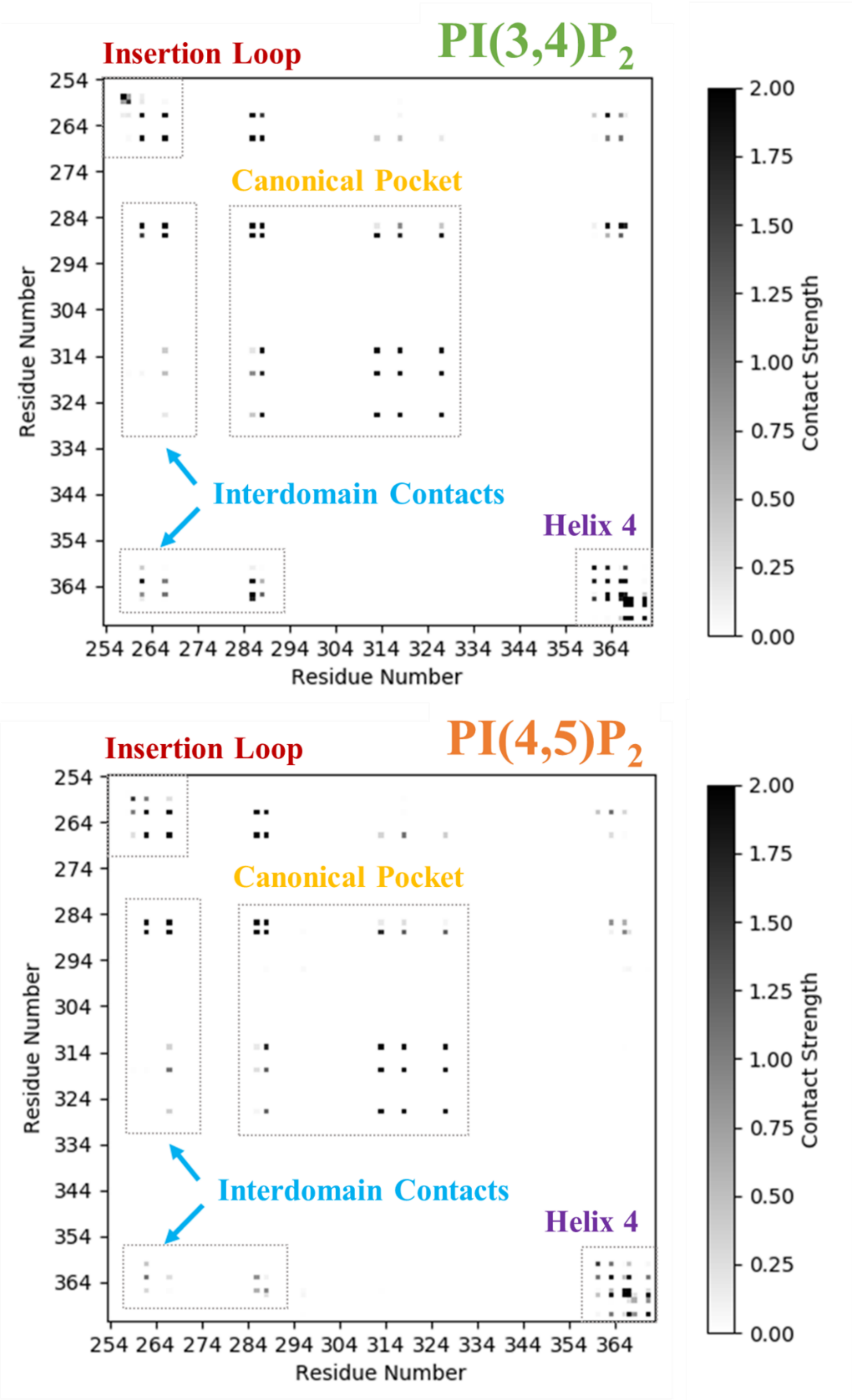


**Fig. S13.** Full PX domain $PIP_2$ contact networks, with PI(3,4)$P_2$ (*Top*) and PI(4,5)$P_2$ (*Bottom*).

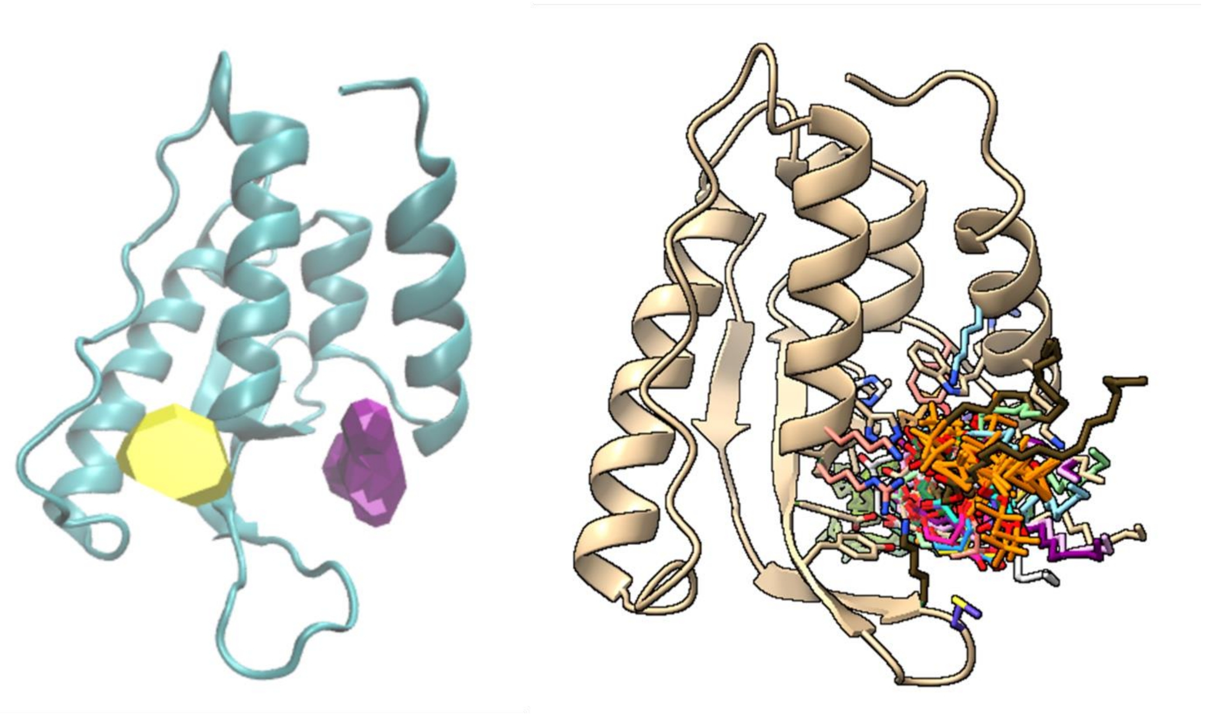

**Fig. S14.** Cavity analysis of the PX domain of 2RAK (*Left*, (60)) and GRAMM docking of the ten most likely predicted $PIP_2$ positions for PI(3,4)$P_2$ and PI(4,5)$P_2$ (*Right*, (61)).
